# Myoferlin is an essential component of late-stage vRNP trafficking vesicles for enveloped RNA viruses

**DOI:** 10.1101/2024.07.02.601679

**Authors:** Stefano Bonazza, Hannah L Turkington, Hannah L Coutts, Swathi Sukumar, Courtney Hawthorn, Erin M P E Getty, Olivier Touzelet, Judit Barabas, Ultan F Power, David G Courtney

## Abstract

The Rab11a endosomal recycling pathway is exploited by important respiratory RNA viruses such as IAV and RSV, aiding viral egress from the apical surface of polarized epithelial cells. Late in infection, Rab11a-containing vesicles specifically transport viral ribonucleoprotein (vRNP) complexes towards the cell surface before packaging and budding. Rather than employing traditional Rab11a-positive recycling endosomes, virus-infected cells generate remodelled Rab11a-containing vesicles, as observed during IAV infection. Besides Rab11a, no other conserved host co-factors have been identified among these various vRNP trafficking vesicles. Here we discovered and confirmed myoferlin’s association with IAV vRNPs in the cytoplasm and colocalisation with Rab11a during late stages of infection. We also found that this role was conserved in late-stage vRNP trafficking of other viruses, including RSV and SeV. Myoferlin likely recruits the EHD family of proteins, which are involved in endosomal biogenesis, to these unique vRNP trafficking endosomes, highlighting myoferlin’s pivotal role in viral replication.

## Introduction

Cumulatively, respiratory viral pathogens including influenza A virus (IAV) and respiratory syncytial virus (RSV), represent a major annual global burden, resulting in the estimated death of around 5 million people every year.^1^ While these two highlighted pathogens are both RNA viruses infecting and transmitting from the human respiratory tract, biologically they are clearly distinct in how they replicate within host cells. For instance, IAV has a segmented negative-sense genome and replicates in the nucleus, while RSV has a single-segment negative-sense genome and replicates in the cytoplasm. Despite these differences, a small number of cellular systems have been found to be widely exploited by these and several other RNA viruses. Rab11a-mediated vesicular trafficking is one such system.^2–6^

The Rab11a-containing endosomal recycling pathway constitutes a major cellular trafficking system widely remodelled by enveloped respiratory RNA viruses.^7,8^ This is true for human pathogens, including IAV and RSV, as well as mammalian respiratory pathogens, such as Sendai virus (SeV).^9^ It has also been speculated that coronaviruses exploit this pathway during egress.^10^ At late stages of infection of each virus, Rab11a-containing vesicles are exploited to traffic viral ribonucleoprotein (vRNP) complexes towards the apical surface so that virions are dispersed into the respiratory airways.^3,11–13^ A better understanding of this cellular trafficking system in both the presence and absence of infection could reveal important, novel insights broadly applicable to many virus families. Additionally, it is now clear that it is not the traditional Rab11a recycling endosomes that are simply hijacked and repurposed for vRNP trafficking.^7,14^ Instead, uniquely remodelled Rab11a-containing irregularly coated vesicles (ICV) appear to form in virus-infected cells, at least for IAV vRNP trafficking, with the involvement of structures from the endoplasmic reticulum.^7,15,16^ Aside from the common utilisation of Rab11a, very little is known of the other co-factors that are likely conserved among these vRNP trafficking vesicles that arise during various viral infections. Therefore, the identification of a co-factor that, during infection by various viruses, is found to associate on newly formed Rab11a-containing vRNP trafficking vesicles would be critical in furthering our understanding of generalised directional vRNP egress.

To this end, an immunoprecipitation-mass spectrometry approach was employed using an epitope-tagged influenza polymerase protein at a late stage of infection to uncover host factors of the viral trafficking endosome, aside from Rab11a, that are essential for influenza virus replication. After analysis of these data, the transmembrane protein myoferlin, MYOF, was found to clearly associate with IAV vRNPs in the cytoplasm at late stages of infection and strongly colocalised with Rab11a in both the presence and absence of infection. Additionally, the replication capabilities of clinical and lab isolates of IAV were strongly reduced after siRNA-mediated knockdown of MYOF expression.

Further investigations were undertaken to establish whether this strong association between MYOF and Rab11a-trafficking vesicles persisted on vRNP trafficking endosomes of other respiratory viruses that traffic with directionality to the apical surface of airway epithelial cells. RSV and SeV were both found to be sensitive to MYOF levels in the cell, in the same manner as observed for IAV, while imaging confirmed a strong association between MYOF, Rab11a and vRNPs on trafficking endosomes, recapitulating the observation made during IAV vRNP trafficking. Finally, it has previously been shown that MYOF can associate with EHD proteins,^17^ which are involved in the late-stage maturation and budding of a subset of endosomes.^18,19^ Therefore, the possibility remained that MYOF in these vesicles recruited EHD proteins as an important step in vesicular biogenesis. On investigation, it was revealed that EHD1 and EHD2 are clearly present within these IAV vRNP trafficking endosomes. Herein, we describe the discovery of MYOF as an essential factor for vRNP trafficking vesicle biogenesis and that it likely mediates the recruitment of EHD proteins to complete vesicular remodelling.

## Results

### Identification of the PA interactome

IAV vRNPs are known to traffic on Rab11a-containing vesicles towards the plasma membrane at late stages of infection. This work aimed to uncover potential host factors, in addition to Rab11a, that are essential for this late stage of virus replication. A reporter strain of influenza A/WSN/33 encoding a FLAG-tagged PA subunit (henceforth referred to as WSN_PA-FLAG) was designed and rescued and was used to selectively immunoprecipitate (IP) the PA interactome as a proxy of the vRNP. Paraformaldehyde (PFA) crosslinking was employed to capture even transient and long-range interactors, while also allowing for the use of more stringent wash steps. Moreover, the IP was carried out in the presence of RNases to focus this search exclusively on the protein-protein interactome of the vRNP (Figure 1A). The specificity and stringency of the IP protocol were assayed by western blotting and silver staining of the input samples and the IP fractions. As a baseline and negative control for the IP, the wild-type influenza A/WSN/33 (WSN_WT) was used. A robust enrichment of PA-FLAG, but not wild-type PA, was observed in IP samples, as well as the absence of actin, which is not a known interactor of the vRNP (Figure 1B). Silver staining demonstrated a clear enrichment in overall proteins after FLAG IP from cells infected with WSN_PA-FLAG over WSN_WT (Figure 1C).

**Figure 1-.**
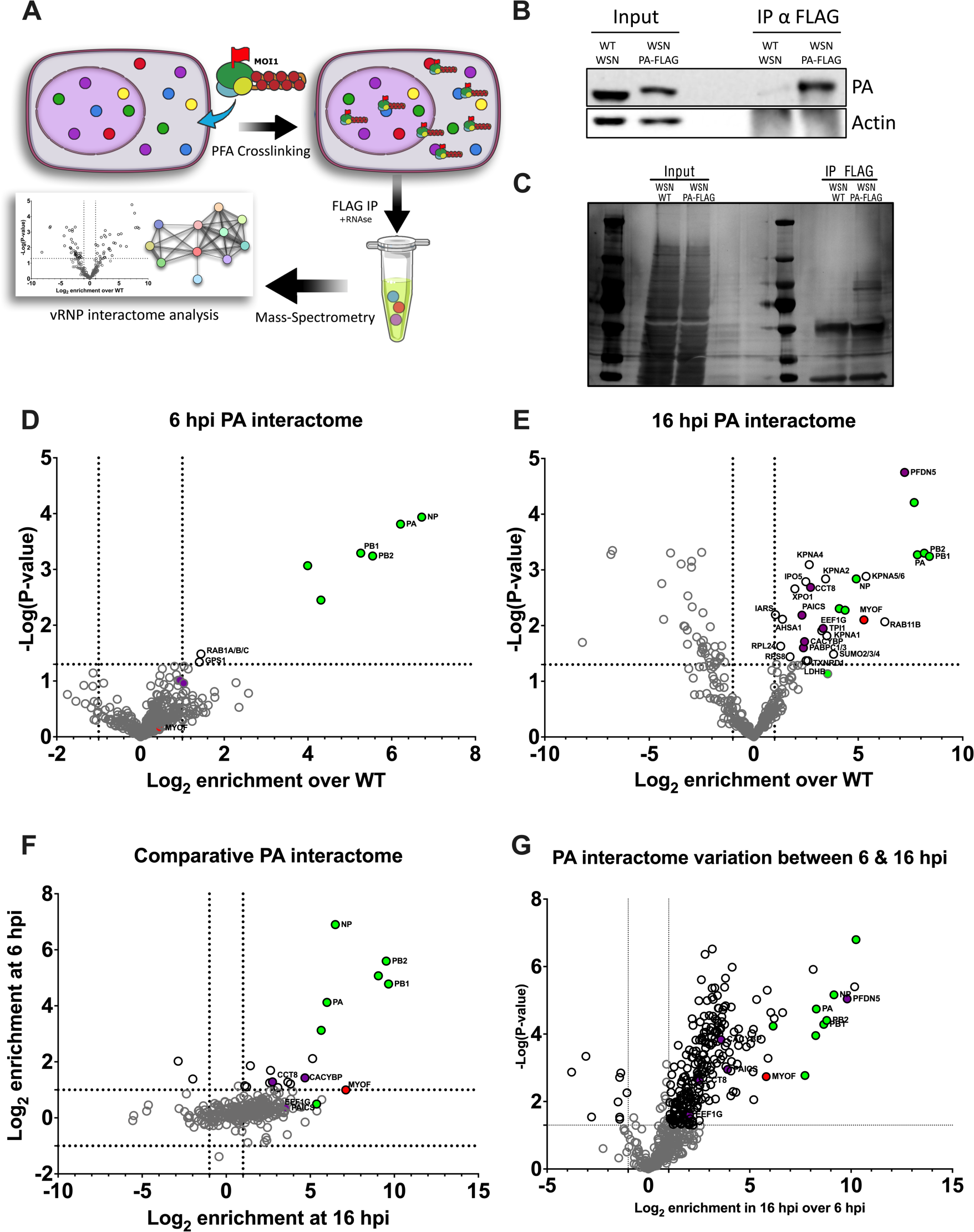
Identification of the PA interactome. (A) Schematic of the immunoprecipitation approach used. A549 cells were infected with the IAV WSN strain encoding a FLAG-tagged PA polymerase subunit for either 16 or 6 h at an MOI of 1, then crosslinked with PFA and harvested. The PA interactome was isolated by FLAG immunoprecipitation in the presence of RNases to isolate exclusively protein-mediated interactions and subjected to mass spectrometry. (B) Representative Western blotting of the 16 hpi IP samples analysed. (C) Representative silver staining of the same IP samples. (D) Volcano plot of the early interactome (6hpi). Viral proteins are represented in green, while purple indicates the hits that were followed up on. Lastly, red indicates myoferlin. (E) Volcano plot of the late interactome (16hpi). (F) Correlation plot of the early and late interactomes. The two datasets were analysed concurrently, and for each entry, an enrichment value for both 6h and 16h was determined. Proteins in the top right quadrant are enriched in both datasets, the ones in the top left are only enriched in the early. (G) Volcano plot representing the differential enrichment of proteins between the early FLAG samples and late FLAG samples. Each dataset is derived from three independent biological replicates.

The experiment was performed at both 6 and 16 hours post-infection (hpi), modelling “early” and “late” infection, respectively, thereby providing a more exhaustive picture of the vRNP interactome throughout the viral replication cycle. The early interactome produced fewer significant hits (Figure 1D; Table S1), perhaps due to the comparatively low expression of viral proteins. In contrast, the late interactome contained proteins involved in translation, protein folding and intracellular protein transport (Figures 1E & S1; Table S2). When comparing proteins enriched in both early and late timepoints, a strong enrichment of the influenza polymerase components PB2, PB1, PA and NP was observed as expected (Figure 1F; Table S3), in addition to host factors including MYOF, CCT8 and CACYBP. Finally, when directly comparing the early to late interactome to assess the dynamics of some of these host interactors, 10 proteins were significantly enriched at 6 hpi vs 16 hpi (Figure 1G, left; Table S4), while a large set of 255 proteins were significantly enriched at 16 hpi vs 6 hpi (Figure 1G, right; Table S4). Interestingly, upon STRING analysis of a subset (enrichment >3 fold) of these 16 hpi enriched proteins, a clustering of proteins associated with the nucleocytoplasmic transport complex was observed, as expected for mature vRNPs shuttling from the nucleus at late stages of infection (Figure S2, circled red). More intriguingly, we found that 14 of these strongly enriched PA-interacting proteins (Figure S2, circled blue) were previously identified as host proteins reliably packaged into WSN virions.^20^ In fact, these 14 proteins make up 40% of the 35 total host proteins identified in this study, and constitute a reliable indicator that this 16 hpi interactome is indeed capturing late-stage vRNP interacting proteins.

### MYOF associates with vRNPs at late stages of infection

These interactomes constitute an important snapshot of host-pathogen interactions. In particular, the late interactome comprised a collection of putative host trafficking determinants. From this dataset, the six most significantly enriched hits were investigated via siRNA knockdown, followed by infection, and quantification of viral replication. A549 cells were transfected with siRNAs targeting CACYBP, CCT8, EEIF1G, MYOF, PAICS, and PFDN5, as well as a non-targeting control (NSC) and PABPC1 as a positive control.^21^ After 48 hours, cells were infected with a reporter A/WSN/33 strain encoding a fluorescently tagged PA subunit (WSN_PA-mNeon). Supernatants were collected at 48 hpi and infectious virus was titred. Knockdown of most candidate genes significantly reduced IAV titres, with only knockdown of PAICS having no effect (Figure 2A). Interestingly, CACYBP, CCT8, MYOF and PFDN5 reduced IAV titres below the limit of detection. Among the candidates that strongly affected IAV titres, MYOF was chosen as a promising candidate for mediating viral trafficking, as previous work linked it to membrane dynamics and endocytosis.^22^

**Figure 2-.**
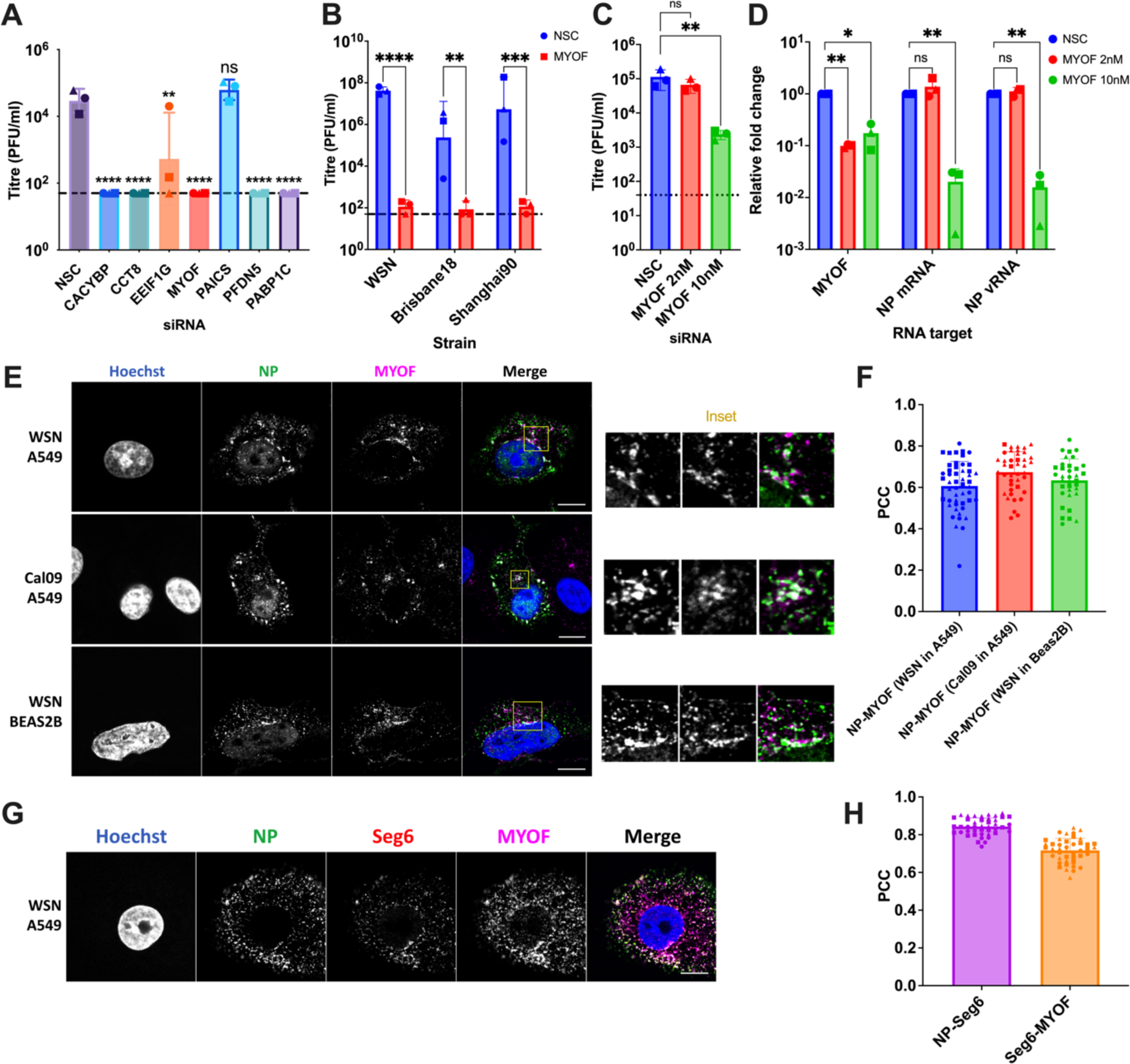
MYOF associates with vRNPs at late stages of infection. (A) Titres of supernatants from siRNA transfected A549 cells that were subsequently infected with MOI 0.01 with the reporter WSN_PA-mNeon for 48h. 20nM of siRNAs were used targeting either a non-specific control (NSC), a positive control (PABPC1) or a candidate hit from the interactome analysis (CACYBP, CCT8, EEIF1G, MYOF, PAICS, PFDN5) (B) Titres from supernatants of siRNA-transfected A549 cells infected at MOI 0.01 with WSN or 0.1 with A/Brisbane/02/2018 (Brisbane18) or A/Shanghai/24/90 (Shanghai90) for 48h. (C) Titres from supernatants of siRNA-transfected A549 cells infected at MOI 5 with WSN for 8h. Two different concentrations (2nM or 10nM) of MYOF siRNA were tested (D) The cells from (C) were then washed and RNA was harvested for RT-qPCR. (E) A549 cells (top and middle row) or BEAS-2B cells (bottom row) were infected with MOI 5 WSN for 8h (top and bottom row) or Cal09 MOI 1 for 16h (middle row), then fixed and stained for NP (green), and MYOF (magenta), and counterstained with Hoechst. The perinuclear accumulation in the yellow inset is magnified on the right. Scale bar 10 µm. (F) Pearson’s correlation coefficient quantification of the colocalisation between MYOF and NP in single cells in the indicated conditions. (G) A549 cells were infected for 8h with MOI 5 WSN, then fixed and processed for smiFISH to stain segment 6 (NA, red), followed by immunofluorescence to visualise NP (green) and MYOF (magenta). Scale bar 10 µm. (H) Pearson’s correlation coefficient quantification of the pairwise colocalisation between NP, MYOF, and segment 6, in single cells. In (F) and (H) different shaped points indicate individual cells from different replicates. All experiments consist of 3 biological replicates and error bars represent standard deviation.

The effect of myoferlin knockdown on IAV was not a strain-specific phenotype, since a similar reduction in replication was observed for the H1N1 lab strain A/WSN/33 (WSN), the post-pandemic H1N1 A/Brisbane/02/2018 (Brisbane18), and the H3N2 strain A/Shanghai/24/90 (Shanghai90) (Figure 2B).

Figure 2C shows the effect of MYOF knockdown on viral titres. Two different concentrations of MYOF siRNA were used to investigate dose-dependent phenotypes. Viral RNA and myoferlin mRNA levels were then quantified at these siRNA concentrations to further assess these differences (Figure 2D). Indeed, although the lower concentration (2nM) seemed to reduce MYOF mRNAs to a similar level to the higher concentration (10nM) (∼10%), only the latter had a significant impact on NP mRNA and vRNA at 8 hpi. It is likely that protein levels are not equivalent at these siRNA concentrations, but these data indicate that IAV replication kinetics may be quite sensitive to low levels of myoferlin expression.

Immunofluorescence was then employed to assess myoferlin’s potential as a trafficking factor for IAV vRNPs. A549 or the immortalised bronchial epithelial cell line BEAS-2Bs were infected with either WSN_WT for 8h or the H1N1 A/California/07/2009 (Cal09) for 16h, then fixed and stained for MYOF and NP proteins, as a proxy for vRNPs. As shown in Figure 2E and quantified in Figure 2F by Pairwise Pearson’s Correlation Coefficients (PCC), myoferlin colocalised extensively with NP in the cytoplasm, particularly in the juxtanuclear region, as shown in the magnified images (inset). Myoferlin’s colocalisation with the vRNPs was further assayed by single-molecule inexpensive fluorescent in situ hybridisation (smiFISH). A549 cells were infected at MOI 5 with WSN_WT for 8h, fixed and stained for NA vRNA (Seg 6), NP protein, and MYOF (Figure 2G). Correlation analysis (Figure 2H) confirmed the strong colocalisation between MYOF and vRNPs.

The significant effect of myoferlin depletion on IAV replication, its extensive colocalisation with vRNPs during late-stage infection, and particularly its intracellular localisation appearing comparable to that of the canonical endocytic recycling machinery, provided a rationale for a deeper study of myoferlin’s role in the recycling pathway alongside Rab11a.

### Rab11a and MYOF work in tandem to coordinate slow endocytic recycling

The potential relationship between myoferlin and Rab11a was initially investigated in the absence of infection, to better ascertain the role myoferlin may play in cellular recycling under normal growth conditions. First, it was observed that MYOF and Rab11a strongly colocalised by immunofluorescence in an A549_Rab11a-mCherry reporter cell line, as shown in Figure 3A and quantified in Figure 3B. Colocalisation was not limited to the prominent juxtanuclear accumulation but extended to the smaller foci at the cellular periphery (as shown in the magnification), suggesting an overlap throughout trafficking.

**Figure 3-.**
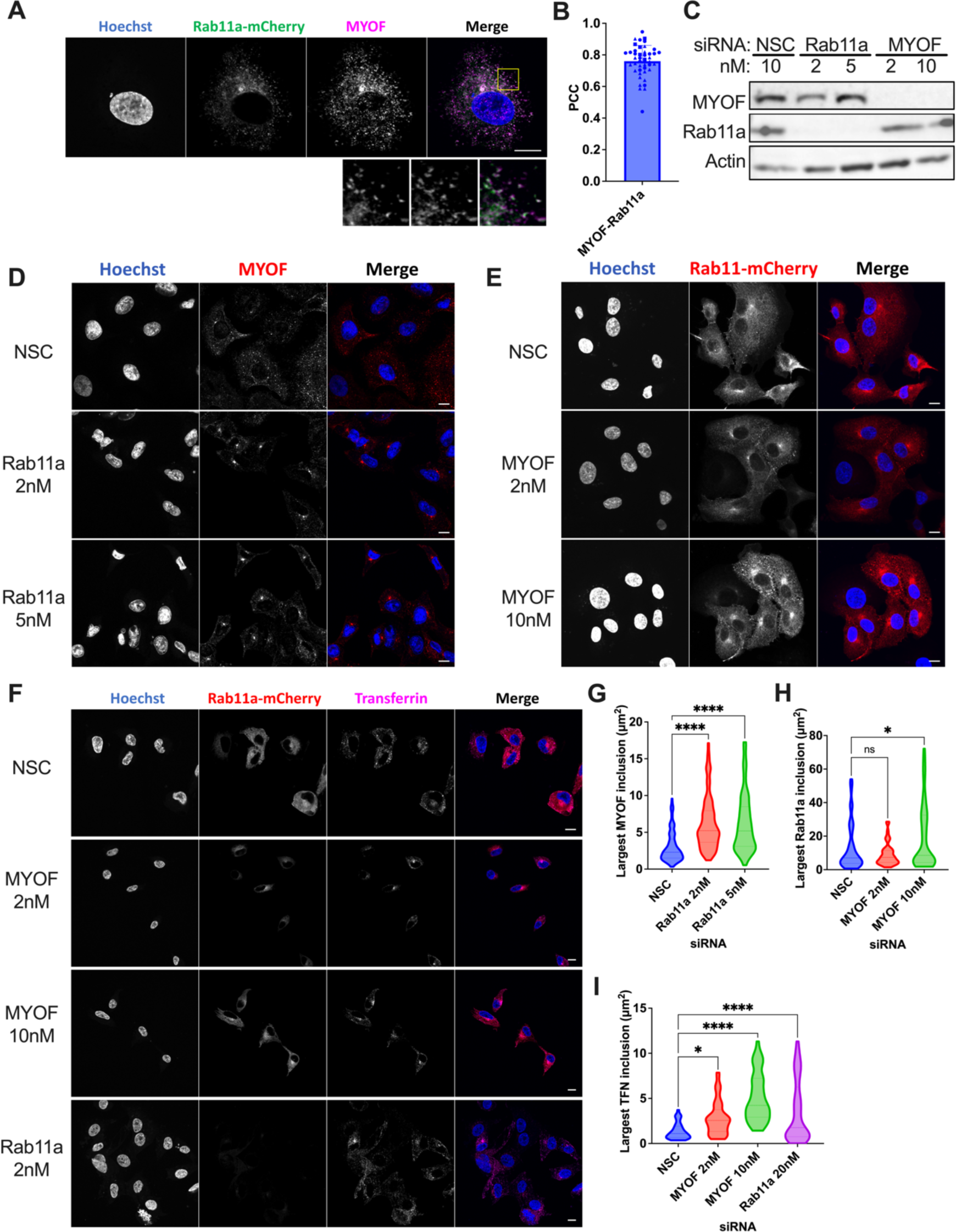
Rab11a and MYOF work in tandem to coordinate slow endocytic recycling. (A) A549_Rab11a-mCherry cells were stained for MYOF and imaged. The yellow inset is shown magnified below. Scale bar 10 µm. (B) Pearson’s correlation coefficient analysis between MYOF and Rab11a in single cells, with different shaped points indicating individual cells from different replicates. (C) Western blot of protein expression in siRNA-treated A549 cells 48h post transfection. (D) siRNA-treated A549 cells stained for MYOF 48h post-transfection. Maximum intensity Z-stack projections are shown. Scale bar 10 µm. (E) siRNA treated A549_Rab11a-mCherry cells imaged for Rab11a-mCherry 48h post transfection. Maximum intensity Z-stack projections are shown. Scale bar 10 µm. (F) siRNA treated A549_Rab11a-mCherry cells transfected for 48h before cells were serum starved for 30’ at 4°C, incubated at 37°C with Atto647-labelled transferrin (Tfn) for 30’ and immediately fixed on ice. Scale bar 10 µm. (G-I) Measurement of the largest signal (G, MYOF; H, Rab11a; I, Tfn) accumulation in each cell, measured by automated thresholding based on image entropy. All experiments consist of 3 biological replicates and error bars represent standard deviation.

The extensive colocalisation suggested the possibility of a functional relationship between MYOF and Rab11a in the orchestration of the endocytic recycling compartment (ERC). Therefore, the effect of the ablation of one protein on the localisation of the other was next explored. The siRNAs successfully reduced myoferlin and Rab11a protein to undetectable levels (Figure 3C). Knockdown of Rab11a in wild-type A549 cells profoundly affected MYOF localisation, with its signal aggregating at a single juxtanuclear spot in the absence of the GTPase (Figure 3D). Similarly, myoferlin depletion led to a pronounced remodelling of the Rab11a network, resulting in larger juxtanuclear accumulations of Rab11a (Figure 3E). As Rab11a localisation is perturbed in the absence of MYOF, it stands to reason that Rab11a-dependent recycling would be altered as well. Indeed, cells depleted for MYOF showed a marked redistribution of Atto647-tagged transferrin in a transferrin uptake assay (Figure 3F). After 30 minutes of transferrin uptake, most of the signal in MYOF-depleted cells accumulated in large foci near the nuclear membrane, while in control cells it was dispersed in smaller distinct puncta throughout the cytoplasm. This was also recapitulated by Rab11a knockdown. These aggregating phenotypes were confirmed by quantifying the area of the largest cluster of signals in the cytoplasm of each cell for each condition (Figure 3G, 3H and 3I).

Taken together, these data suggest a role for myoferlin in Rab11a-mediated recycling in uninfected cells, thereby confirming (at least in this A549 lung-cell model) the intimate relationship between the two. However, during IAV infection the Rab11a+ ERC is severely remodelled, affecting the localisation, composition, and recycling function of the whole vesicular network.^7,15^ Therefore, the presence of MYOF in these distinct virus-induced vesicles was next explored.

### MYOF is specifically retained in influenza A vRNP-trafficking vesicles

IAV infection profoundly affects the ERC, particularly during late-stage infection. It remodels the structure and composition of Rab11a+ vesicles. So far, we have shown that Myoferlin and Rab11a work in concert to ensure proper recycling in healthy cells, but nothing is known about myoferlin’s fate upon infection with IAV. To adequately observe the remodelling of the ERC upon infection, the reporter A549_Rab11a-mCherry cell line was infected for 8h with WSN_WT and imaged for NP, mCherry and MYOF. Compared to mock-infected cells, infected cells showed a marked redistribution of the Rab11a-mCherry signal, displaying several enlarged foci strongly colocalising with NP, and importantly MYOF (Figure 4A and 4B).

**Figure 4-.**
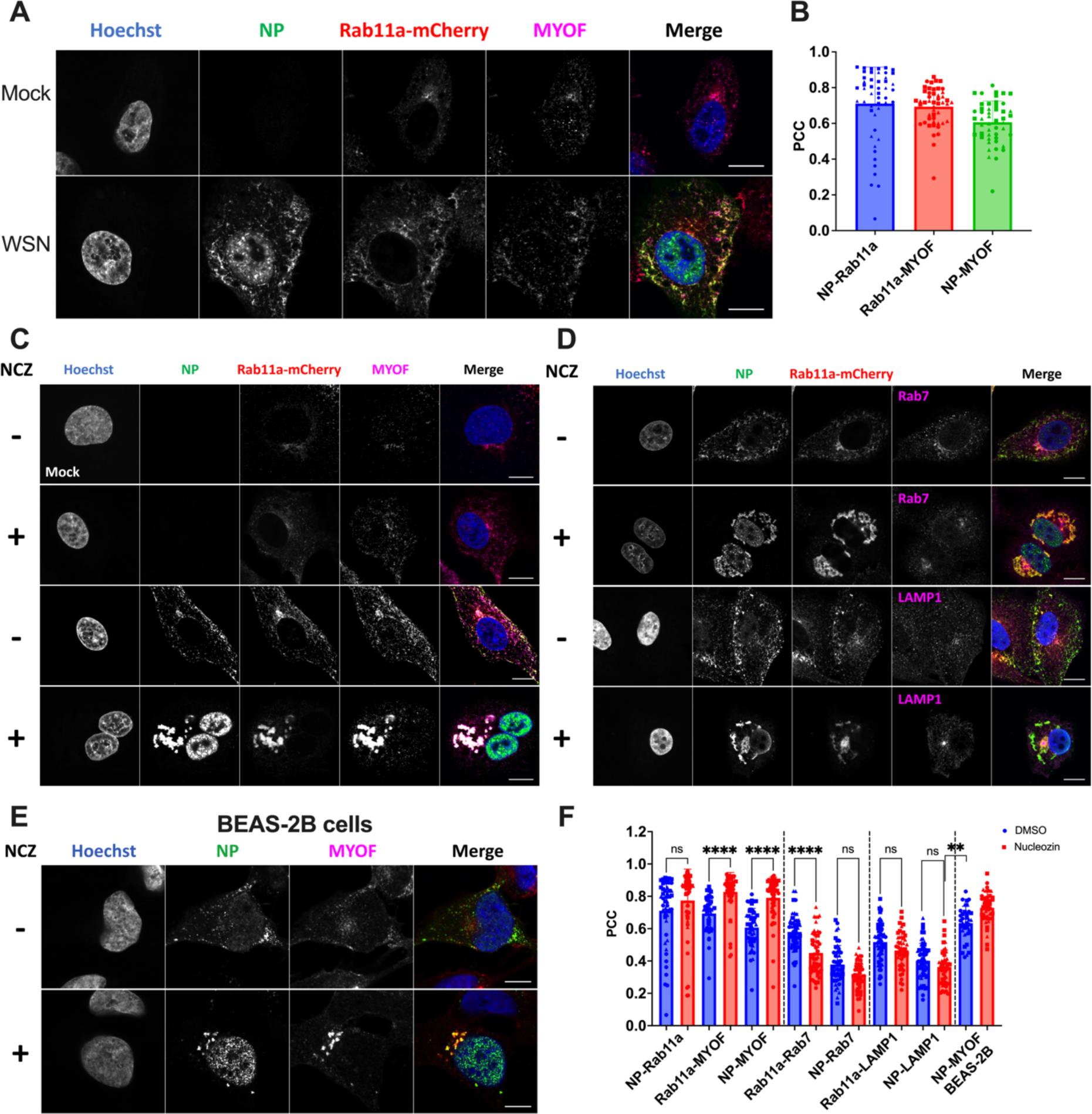
MYOF is specifically retained in influenza A vRNP-trafficking vesicles. (A) A549_Rab11a-mCherry cells were mock-infected or infected with MOI 5 WSN for 8h, then fixed and stained for NP (green) and MYOF (magenta). Scale bar 10 µm. (B) Pearson’s correlation coefficient quantification of the pairwise colocalisation between NP, MYOF, and Rab11a, in single cells. (C-D) A549_Rab11a-mCherry cells were infected with MOI 5 WSN for 6h, treated or mock-treated with nucleozin (NCZ) for 2h, then fixed and stained for NP and MYOF (C), or NP and either Rab7 or LAMP1 (D). Scale bar 10 µm. (E) BEAS-2B cells were infected with MOI 5 WSN for 6h, treated or mock-treated with nucleozin (NCZ) for 2h, then fixed and stained for NP and MYOF. Scale bar 10 µm. (F) Single-cell Pearson’s correlation coefficient quantification of colocalisation between NP and MYOF, or NP and the controls (Rab7/LAMP1). In blue are represented the mock-treated cells, and in red the nucleozin-treated. In (B) and (F) different shaped points indicate individual cells from different replicates. All experiments consist of 3 biological replicates and error bars represent standard deviation.

Colocalisation analysis on its own would not be sufficient to prove MYOF’s inclusion in irregularly coated vesicles (ICVs), due to the small and dispersed nature of these vesicles. To bypass this issue, infected cells were treated at 6hpi with nucleozin, an NP-crosslinking drug,^23^ to induce the selective aggregation of vRNP-associated vesicles. Thus, ICVs and all their constituents would coalesce into easily detectable foci. As shown in Figure 4C, the drug had no effect in the absence of infection but caused the formation of large inclusions in infected cells. As expected, these inclusions were positive for NP and Rab11a. Interestingly, they were also MYOF-positive. Importantly, nucleozin addition did not alter the localisation of other subcellular compartment markers such as Rab7 (late endosomes) or LAMP1 (lysosomes) (Figure 4D), indicating a specific effect on vRNP trafficking vesicles only. Moreover, these observations were confirmed in BEAS-2B (Figure 4E). PCCs were calculated for single cells in the presence or absence of the drug, showing an increased colocalisation of NP and MYOF, NP and Rab11a, and Rab11a and MYOF, but not for the negative controls Rab7 and LAMP1, following aggregation (Figure 4F). This established the presence of MYOF in ICVs, providing evidence of its likely functional partnership with Rab11a even during infection.

### Respiratory viruses commonly utilise Rab11a-MYOF vesicles at late stages of infection

The trafficking of vRNP complexes of respiratory viruses other than IAV, including RSV and SeV, is known to depend on Rab11a-containing vesicles. To investigate whether replication of these viruses is also dependent on myoferlin, smiFISH imaging was performed to visualise RSV or SeV vRNAs during infection of A549 cells. These data revealed a high degree of colocalisation between the genomic RNAs and MYOF (Figures 5A and 5C). A549_Rab11a-mCherry cells were infected with RSV_BT2a (a low passage clinical isolate), while A549 cells were infected with SeV_eGFP. After 72 h the samples were stained for their respective vRNAs and MYOF. For both viruses, the most abundant colocalisation occurred within smaller, irregularly shaped vRNA aggregates and not within larger replication factories. The same phenomenon was also observed for a different RSV strain (RSV_mKate2, Figure S4). This is consistent with a model in which the Rab11a-MYOF pathway is used specifically to transport mature genomes towards sites of virion budding. Interestingly, for RSV-infected cells, infection seemed to induce a strong re-localisation of the MYOF signal, indicative of ERC remodelling. At least for RSV-infected cells, MYOF staining colocalised well with the Rab11a-mCherry signal, confirming their relationship even during RSV infection. Furthermore, depletion of MYOF could not support effective replication of RSV and SeV, recapitulating Rab11a depletion (Figure 5B and 5D).

**Figure 5-.**
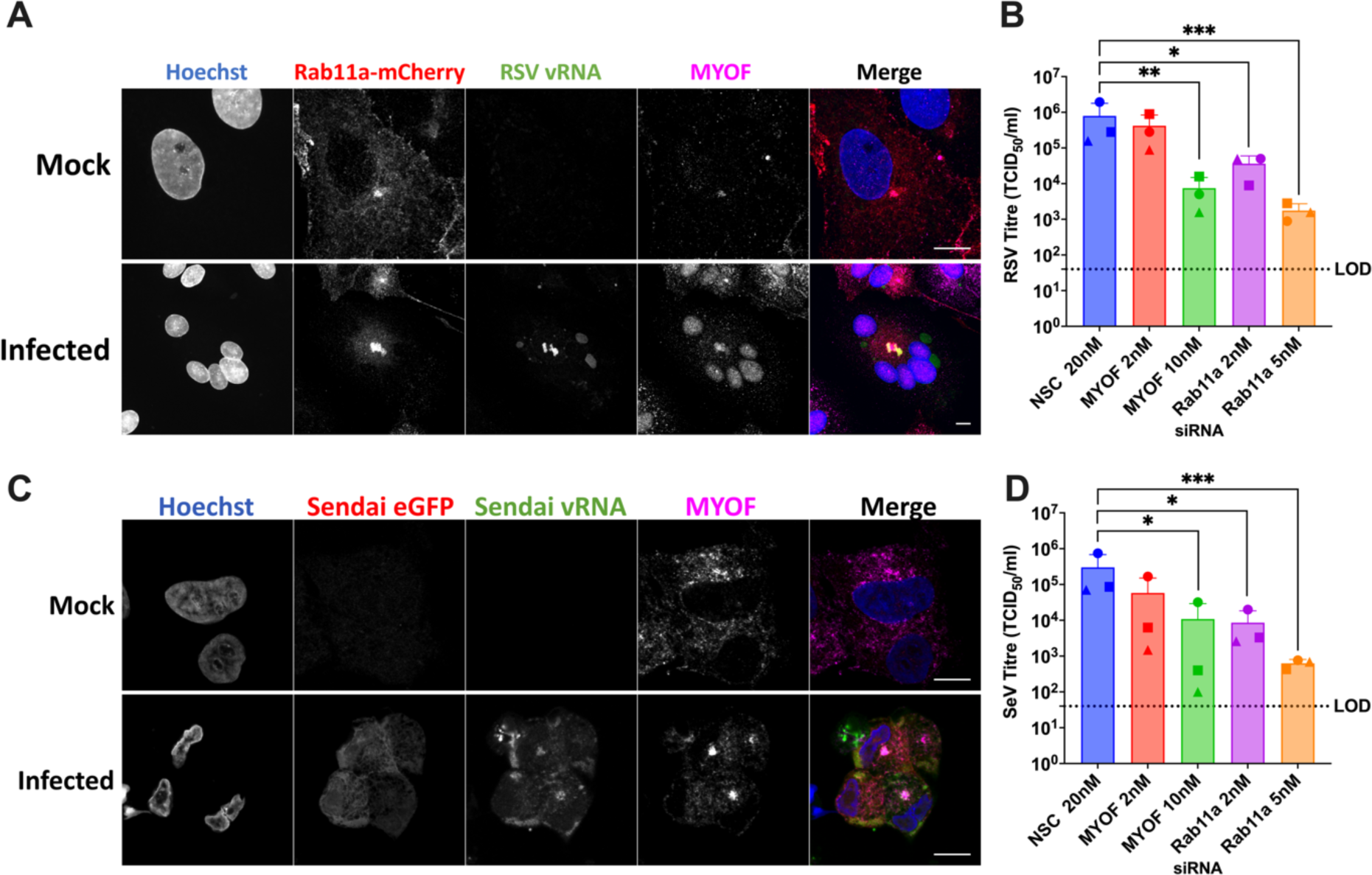
Respiratory viruses commonly utilise Rab11a-MYOF vesicles at late stages of infection. (A) A549_Rab11a-mCherry cells were mock-infected or infected with RSV_BT2a for 72h, then processed for smiFISH to image the viral genomes (green), as well as immunofluorescence for MYOF (magenta). Scale bar 10 µm. (B) A549 cells were transfected with the indicated siRNA concentrations for 48h, then infected with RSV_mKate2 for 72h. Supernatants were harvested and titred. (C) A549 cells were mock-infected or infected with Sendai_eGFP for 72h, then processed for smiFISH to image the viral genomes (green), as well as immunofluorescence for MYOF (magenta). Scale bar 10 µm. (D) A549 cells were transfected with the indicated siRNA concentrations for 48h, then infected with Sendai_eGFP for 72h. Supernatants were harvested and titred. All experiments consist of 3 biological replicates and error bars represent standard deviation.

These data confirm myoferlin as an active player in vRNP trafficking along the Rab11a recycling pathway for multiple enveloped viruses. As such, they provide novel insights into multi-species enveloped virus vRNA trafficking in respiratory cells and, by extension, intriguing targets for the design of broad-spectrum antivirals.

### MYOF likely recruits EHD1/2 for end-stage ICV biogenesis

Myoferlin is a large protein composed of several C2 domains, some of which were characterised as calcium binding, membrane binding, or protein binding. Specifically, the second C2 domain (C2B) was found to be responsible for direct interaction with Eps15 Homology Domain 2 (EHD2).^17^ To assess whether EHD2, or the closely related EHD1, are present within these ICVs, A549_Rab11a-mCherry cells were transduced with lentiviral vectors expressing GFP-tagged forms of the protein of interest, or a negative control GFP-only (EV). They were then infected with WSN_WT for 6h, treated with nucleozin for 2 h, and imaged. In untreated cells, both EHD proteins localised to distinct cytosolic structures, including the perinuclear area where they partially colocalised with Rab11a, and therefore likely myoferlin (Figure 6A). In contrast, nucleozin treatment generated large and easily recognisable Rab11a aggregates, with both EHD1 and EHD2 predominantly localising therein (Figure 6A). These data confirm the presence of members of the EHD family of proteins, specifically EHD1 and EHD2, in ICVs. Given their known role in vesicular biogenesis and the association between EHD2 and MYOF,^17^ we propose that during infection with respiratory viruses, such as IAV, RSV, or SeV, EHD proteins are recruited to vRNP trafficking vesicles by MYOF to coordinate membrane remodelling and complete ICV maturation (Figure 6B). This would lead to trafficking of vRNPs to the cytoplasmic membrane for packaging into virions and subsequent egress.

**Figure 6-.**
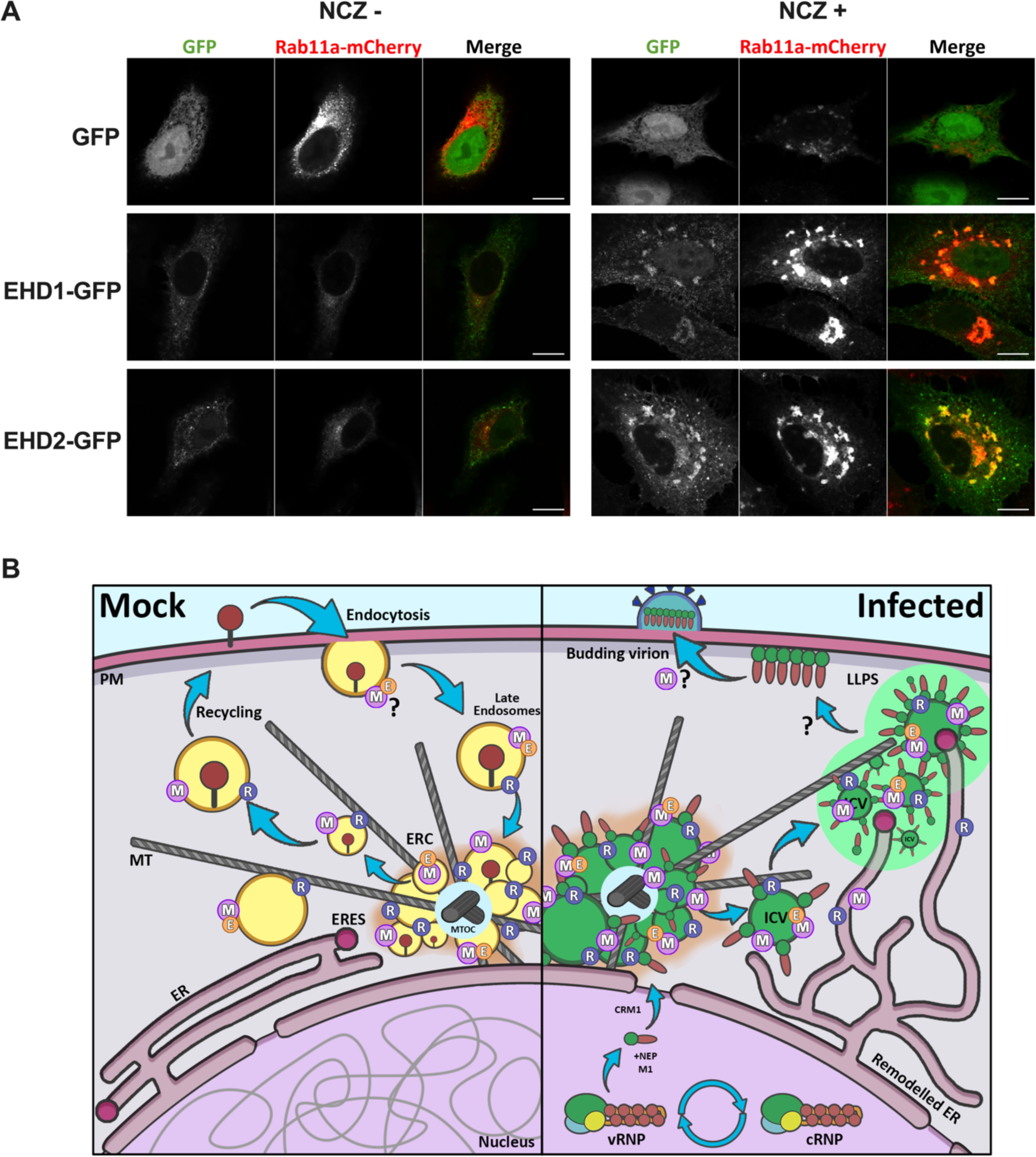
MYOF likely recruits EHD1/2 to complete vRNP-trafficking vesicle biogenesis. (A) A549_Rab11a-mCherry cells were transduced with a vector expressing either GFP, EHD1-GFP or EHD2-GFP. They were then infected with MOI 5 WSN for 6h and treated (left) or mock-treated (right) with nucleozin (NCZ) for 2h before fixation and imaging. Scale bar 10 µm. Images are representative of 3 biological replicates. (B) Schematic showing MYOF and EHD1/2 incorporation into vRNP-trafficking ICVs. On the left, in uninfected cells MYOF associates with Rab11a recycling endosomes at the ERC and is actively involved in the transport of cellular components including receptors and transferrin for recycling. On the right, in IAV infected cells the ER is remodelled while vRNPs exported from the nucleus congregate at the MTOC. These vRNPs are trafficked to the cellular membrane for packaging on ICVs containing Rab11a and MYOF, with vesicular maturation possibly triggered by the recruitment of EHD1 and EHD2. ICVs are proposed to traffic to the ERES where vRNPs may then undergo LLPS prior to packaging into virions. PM: Plasma Membrane, MT: Microtubules, ERC: Endosomal Recycling Compartment, ER: Endoplasmic Reticulum, ERES: ER Egress Sites, MTOC: MT Organising Centre, M: Myoferlin, R: Rab11a, E: EHD1/2, LLPS: Liquid-Liquid Phase Separation, ICV: Irregularly Coated Vesicles.

## Discussion

Due to their relatively small genome sizes, and thus limited coding capacity, RNA viruses rely on host machinery to accomplish most, if not all, of the steps in their replication cycle. From the moment of infection, a series of intricate pathways are exploited and hijacked to ensure viral replication. IAV intracellular trafficking is very well-studied and an ideal model to investigate the dynamic movements of viral RNA genomes. Firstly, upon IAV virion binding to the plasma membrane sialic acid receptors, clathrin-mediated endocytosis is triggered (with a minority of particles entering through other means) and the virion is trafficked in the endosomal pathway.^7^ Acidification of the compartment induces the release of vRNPs in the cytosol, with nuclear localisation signals on the vRNP recruiting importins to carry it across the nuclear pore.^24^ After replication in the nucleus, newly formed vRNPs interact with M1 and NEP to hijack the CRM1 (XPO1) pathway for export to the cytosol.^24,25^ Once through the nuclear pore complex, vRNPs rapidly concentrate close to the nuclear membrane in the vicinity of the microtubule organising centre (MTOC), and associate with Rab11a+ vesicles through direct interaction of Rab11a with PB2.^26,27^ Seminal publications from the Naffakh lab and the Amorim lab clearly established the extensive remodelling of the vesicular network upon IAV infection.^7,16,28^ Electron microscopy and live-imaging experiments proved the formation of ICVs, characteristic vRNP-clad Rab11+ vesicles disrupting the normal ERC (Figure 6B). ICVs are generated and trafficked along microtubules (MTs). Interestingly, the microtubule network does not seem to be strictly required for IAV replication, so there must be other factors at play.^29^ ICVs then localise near the endoplasmic reticulum (ER), at ER egress sites (ERES), where they are unloaded from MTs by ATG9A.^7,15,16^ The high local concentration of vRNPs triggers liquid-liquid phase separation (LLPS), in which free vRNPs and ICVs coalesce in a membraneless, liquid-like state. These liquid vRNP droplets are believed to be sites for vRNP reassortment and possibly bundling.^15^ Bundles of vRNPs are then transported to budding sites at the plasma membrane. The mechanisms surrounding this last trafficking step are still unclear, however, vRNPs are thought to bind M1 at the cytosolic face of specialised lipid rafts enriched in cholesterol and viral glycoproteins. Molecular crowding, as well as specific membrane-bending activities of M1, M2, HA and NA, induce budding and scission.^30^ These processes completely overhaul the trafficking networks of the cell, with the involvement of the ERC, the ER, the lysosomal pathway, and the Golgi apparatus. Hence, an important focus of IAV research is on characterising the exact composition of vRNP trafficking vesicles.

To this end, we proceeded to perform an interactome study to investigate the host factors that are essential to the late stages of trafficking for IAV vRNPs (Figure 1A). Through IP-MS at an early (6 hpi; Figure 1D) and late (16 hpi; Figure 1E) timepoint, coupled with PFA crosslinking and RNase digestion, we generated a snapshot of proteins and pathways associating with the influenza polymerase complex (FluPol) (Figure 1F). The FluPol is a dynamic entity that interfaces with a multitude of cellular processes over the course of a single infectious cycle, with our method capturing the polyhedric nature of this. By directly comparing early to late datasets we could then further demarcate the host factors likely contributing to vRNP trafficking, bundling, and packaging (Figure 1G). Previous studies aimed to characterise IAV interactomes, mainly by expression of individual or subsets of viral proteins, or in infected cells without any crosslinking.^31–35^ Our study forgoes the purification of exclusively direct interactors in favour of isolating more indirect, long-range interactions still fundamental for viral replication, in the context of the remodelled infected cell.

The quantification of proteins enriched in the late sample compared to controls, identified groups of proteins associated with intracellular protein transport, translation and protein folding (Figure S1). This is likely due to the active translation and trafficking of mature FLAG-tagged PA, and the nuclear export of vRNP complexes. However, because this method of quantification can only quantify proteins present in both FLAG (WSN_FLAG) and control (WSN_WT) datasets, we were more interested in investigating the proteins enriched in the late compared to early post-infection timepoints. Here we found two clear groups of proteins: those involved in nucleocytoplasmic transport, as expected for exporting vRNP complexes and, most interestingly, 14 proteins previously described to be incorporated into WSN virions and released from the cell. ^20^ This validates our timepoint, clearly indicating the capture of late interactors of the FluPol complex, while substantiating the discovery of distinct packaged host factors into IAV virions. Interestingly, a few hits were strongly enriched in the late over early interactome and had no previously known roles in IAV replication. Our siRNA knockdown and infection assays, coupled with prior literature on its known functions, identified myoferlin as the main host factor of interest.

Myoferlin is a large multidomain protein, part of the ancient family of Ferlin proteins. These transmembrane factors are involved in vesicular trafficking and membrane dynamics in a myriad of contexts.^22,36^ Myoferlin was first characterised as an important factor in muscle cell maturation^37,38^ but has since been linked to a plethora of processes in vastly different cellular environments. Studies have proven myoferlin’s involvement in mitochondrial fission/fusion events,^39^ lysosomal membrane damage repair,^40^ endothelial growth factor receptor function,^41^ excitation/contraction coupling in mouse skeletal muscle,^42^ invasive phenotype in various cancer types,^43^ and endocytosis.^22,44^ In fact, a recent study found that myoferlin ensures proper localisation of the Env protein to sites of virion budding in human T-cell leukaemia virus type 1 (HTLV-1) infection.^45^

In this work, we show that myoferlin is an important component of the Rab11a+ recycling endosomes, colocalising extensively with the GTPase in uninfected lung epithelial cells, influencing its localisation and its function, while MYOF’s localisation is itself affected by Rab11a.

Endosomal recycling is a fundamental host pathway involved in the vesicular trafficking of a myriad of cargo, such as internalised receptors and other endocytosed material.^14^ Under normal growth conditions recycling endosomes spread out in the cytoplasm but are concentrated in a characteristic juxtanuclear accumulation, with myoferlin localisation closely resembling this typical recycling endosome pattern (Figure 3A). Additionally, it was reported that myoferlin-null myoblasts display delays in transferrin turnover in pulse-chase experiments.^17^ Collectively, these data led us to hypothesise that myoferlin is an important component of endocytic recycling in lung epithelial cells, which is usually the target for infection by respiratory viruses such as IAV and RSV. Using A549 cells as a model we discovered that Rab11a and myoferlin are intimately connected through their roles in recycling, with each dependent on the other for normal cellular distribution (Figure 3D & 3E) and transferrin recycling wholly dependent on both (Figure 3F). With the interdependence of these two components established, coupled with the knowledge that Rab11a-mediated trafficking of vRNPs is a critical step in IAV replication, it is clear that the inhibitory effect of myoferlin loss on viral titres may be attributed to this late-stage in the viral replication cycle. Mounting evidence suggests that several RNA viruses exploit similar strategies to accomplish directional intracellular trafficking of their vRNPs.^4,6,8,46^ Given the strong link that we established between Rab11a and MYOF both in uninfected cells and during IAV infection, we reasoned that it was likely that other pathogens that hijack this pathway would also require myoferlin for efficient replication. Indeed, myoferlin knockdown followed by RSV or SeV infection greatly reduced overall infectious virus release (Figure 5B & 5D), mirroring the results observed following IAV infection (Figure 2C). Furthermore, upon imaging of RSV vRNA, Rab11a-mCherry and MYOF, a clear localisation of all three signals to multiple defined cytosolic spots was evident (Figure 5A). Again, this is a phenotype recapitulated in SeV-infected cells (Figure 5C). Interestingly, it appears as though the largest cytosolic spots, likely active viral replication factories, do not colocalise with Rab11a or myoferlin. This implies that both Rab11a and myoferlin are not required for actual transcription or replication, as we would expect, but instead are only recruited to or associate with actively trafficking mature vRNP-containing vesicles. This clearly aligns with our observations and those of other studies investigating the role of Rab11a strictly in vRNP egress. While only three respiratory viruses that utilise Rab11a-mediated vRNP trafficking were investigated here, these data imply a conserved role for myoferlin in the replication of other Rab11-dependent viruses, including respiratory pathogens of the Coronaviridae and Paramyxoviridae families.^3,10,47^

In establishing myoferlin as an important component of recycling in healthy cells and vRNP trafficking in a range of viral infections, we were interested to note that the second C2 domain (C2B) was previously reported to directly interact with EHD2.^17^ The EH domain is a protein-protein interaction domain often associated with vesicular trafficking factors. It functions by binding cognate NPF/KPF motifs in partner proteins. EHD proteins themselves contain both the eponymous EH domain and the NPF/KPF motif, allowing for self-oligomerisation.^48^ The formation of multimeric EHD structures drives membrane remodelling events and contributes to vesicular trafficking. Myoferlin itself contains an NPF/KPF motif in its second C2 domain, which was found to be fundamental for EHD2 interaction.^17^ Therefore, we hypothesised that, with the presence of MYOF in the ERC, but more importantly within ICVs during IAV infection, members of the EHD family of proteins would be recruited to these vesicles to trigger membrane remodelling and end-stage vesicular maturation (Figure 6B). Indeed, as shown in Rab11a-mCherry cells expressing eGFP-tagged EHD1 and EHD2, upon IAV infection both proteins were found to be localised close to Rab11a at the MTOC. This observation was even starker upon nucleozin treatment to aggregate vRNPs and their accompanying interactors (Figure 6A). These findings lead us to propose that the membrane remodelling observed, particularly during IAV infection, may be triggered by EHD proteins after their recruitment to the ERC by myoferlin. Interestingly, it has also been speculated that the membrane remodelling itself could be the driving force behind vRNP movement towards the cell membrane.^7^ However, very little work was carried out to explore the role of EHDs in viral infections, with only limited evidence supporting EHD1 involvement in Vesicular Stomatitis Virus Glycoprotein (VSV-G) localisation to the plasma membrane.^50^

As stated, important research is being carried out to fully characterise the makeup of influenza trafficking vesicles. So far very little progress has been made towards this goal, but this manuscript extends the list by three, adding MYOF, EHD1 and EHD2. All these factors clearly colocalise with ICVs during trafficking and aggregate upon nucleozin treatment. More work is needed to completely elucidate their role in infection, as well as their involvement in the replication cycle of other pathogens.

Interestingly, MYOF ablation also seemed to affect viral mRNA and vRNA production (Figure 2D), in contrast to the established literature around Rab11a and IAV. As discussed above, myoferlin is believed to play a role in endocytosis as well as recycling, and we believe that its importance in the viral replication cycle is not limited to late trafficking. Myoferlin *in silico* interactome mapping (Figure S3) revealed the extent of MYOF’s involvement in membrane processes, interacting with endocytosis, recycling, membrane repair and trafficking factors. Interestingly, one of the main hits of this STRING interactome mapping is Annexin-A1 (ANXA1), which was also among the enriched hits in our late PA interactome. ANXA1 was found to be an important determinant of vRNP trafficking from endosomes to the nucleus,^49^ again connecting MYOF, endocytosis and vRNP trafficking. Future work aims to better characterise MYOF during the earlier steps of infection.

At the other end of the infection cycle, myoferlin was shown to localise to plasma membranes and specifically lipid rafts, where it interacts with Dynein-2 to ensure membrane resealing after fission-based endocytosis.^22^ As stated above, specialised lipid rafts are the sites on the cell surface at which IAV budding occurs.^51^ An attractive hypothesis puts myoferlin at the crossroads of membrane remodelling events throughout the viral replication cycle, from endocytosis to ICV remodelling to budding at the plasma membrane.

Novel, broadly applicable therapeutic interventions provide the greatest cost-to-benefit ratio for biomedical research. One avenue in the design of such a strategy is the discovery and subsequent targeting of a ubiquitous host protein essential during a key stage in the replication cycle of various clinically relevant human viruses. One such seemingly pervasive stage is the egress of viral genomic RNA exploiting the endosomal recycling pathway. Herein, we describe the intimate relationship between Rab11a-containing vesicles and myoferlin, under normal growth conditions and within IAV-induced remodelled ICVs. Importantly, however, this relationship is conserved on RSV and SeV vRNA-trafficking vesicles. While the data discussed here directly present strong evidence for the importance of myoferlin for vRNA trafficking of IAV, RSV and SeV specifically, they imply that this would also be the case for all Rab11a-dependent viruses, including coronaviruses,^10^ parainfluenza viruses,^9^ Hantaviruses,^4^ ebolaviruses^6^ and flaviviruses.^5^

## Acknowledgements

This research was funded in part by an ERC-STG grant, PTFLU 949506 awarded to D.G.C. This research received infrastructure support from the Wellcome-Wolfson Institute for Experimental Medicine at Queen’s University Belfast. We would like to thank Adam McShane and Dr Dessi Malinova for providing the Atto 647N-conjugated transferrin used in the transferrin uptake experiments.

## Author contributions

Conceptualization, S.B., and D.G.C.; Methodology, S.B., H.L.T., H.L.C., S.S., C.H., E.M.P.E.G., O.T., and J.B.; Formal Analysis, S.B., H.L.T., H.L.C., C.H., E.M.P.E.G., O.T., and J.B.; Investigation, S.B., H.L.T., H.L.C., C.H., E.M.P.E.G., and O.T.; Writing – Original Draft, S.B., and D.G.C.; Writing – Review & Editing, S.B., H.L.T., H.L.C., S.S., C.H., E.M.P.E.G., O.T., J.B., U.F.P., and D.G.C.; Supervision, U.F.P., and D.G.C.; Funding Acquisition, D.G.C.

## Declaration of interests

The authors confirm that they have no declarations of interest in relation to this work.

## Supplemental information

Document S1 containing Figures S1, S2, S3 & S4 and Tables S5-S8

Table S1-4 Excel file containing LC-MS/MS datasets too large to fit in a PDF, related to Figure 1.

## Online Methods

### Cells

For this work immortalised A549 (ATCC; CCL-185), HEK 293T (ATCC; CRL-3216), BEAS-2B (ATCC; CRL-3588), Hep-2^52^ and MDCK (CCL-34) cells were used. All cells were cultured in DMEM supplemented with 1% Pen-Strep (Thermo Fisher Scientific; 15140122) and 5% FBS (Thermo Fisher Scientific; 10270106) at 37°C and 5% CO_2_.

### IAV virus stocks and infections

WSN (A/WSN/33) viral stocks were generated from a reverse genetics system that has been described previously.^53^ WSN_PA-mNeon virus was generated from the same reverse genetics system, where the pPolI_PA plasmid was substituted for the pPolI_PA-mNeon plasmid encoding for a C-terminal mNeon followed by a duplicated packaging signal, and has been described previously.^54^ Additionally, a C-terminally FLAG-tagged PA virus was also rescued from the same system, pPolI_PA-FLAG. Cal09 (A/California/7/2009), Shanghai90 (A/Shanghai/24/1990), Brisbane18 (A/Brisbane/02/2018) viruses were grown from isolates acquired from the National Institute for Biological Standards and Control, UK. All stocks were grown on MDCK cells in IAV growth media consisting of DMEM, Pen-Strep, 0.2% BSA (Merck; A8412), 25mM HEPES (Merck; H0887) and TPCK-trypsin (Merck; T8802), and titred on MDCK cells by plaque assay.

### Immunoprecipitation for LC-MS/MS

For the late timepoint influenza interactome experiment, approximately 2*10^7 (4*10^7 for the early interactome) A549 cells per condition were seeded on 15cm plates 24h prior to infection. The next day, cells were infected with IAV WSN_PA-FLAG (FLAG) or A/WSN/33 (WT) at an MOI of 1, then fixed at 16hpi (6hpi for the early interactome) by a 10’ incubation with 1% PFA in PBS. Residual PFA was quenched with a 10’ treatment with 1.25M Glycine in PBS and washed three times with cold PBS. Cells were then scraped and pelleted. The samples were lysed in RIPA buffer supplemented with protease inhibitors (Merck; 4693116001) for 1h on ice. Afterwards, cell debris was removed by centrifugation and 10% of the remaining sample was taken as input.

The immunoprecipitation was carried out using magnetic protein G beads (Thermo Fisher Scientific; 88848), which were washed and incubated with monoclonal anti-FLAG antibodies (Sigma; F1804) for 45’ before use. We employed 100 µl of beads, 50 µl of antibodies, and 10 µl of RNAse Cocktail (Thermo Fisher Scientific; AM2286) per condition. After capturing for at least 2h, the beads were thoroughly washed in buffers of decreasing stringency: RIPA buffer, high salt buffer (5% HEPES-KOH, 50% KCl, 0.05% NP40 in H2O), and PBS; all ice-cold and supplemented with 1% DTT. After removing 10% of the beads for western blotting, the rest was resuspended in H_2_O and sent for mass spectrometry analysis. Three independent biological replicates were produced and analysed for each condition.

### Western blot

Western blotting was performed as previously described.^55^ The antibodies used can be found in the Supplementary Table S5. Protein bands on blots were visualised by chemiluminescence.

### Silver staining

Silver staining was performed according to the manufacturer’s protocol, using the Pierce Silver Stain Kit (Thermo Fisher Scientific; 24612), using samples processed as detailed in the IP-MS paragraph.

### LC-MS/MS and data analysis

Mass spectrometry was performed at the Cambridge Centre for Proteomics. Sample treatment, preparation and trypsin digestion were performed exactly as described elsewhere.^54^ Peptide pools were quantified by a Qubit protein assay (Thermo Fisher Scientific). Data were acquired on an Orbitrap™ Fusion™ Lumos™ Tribrid™ mass spectrometer with an equal amount of each sample sequentially loaded over individual 60-minute runs with no fractionation given the likely low complexity of the sample. Raw mass spectrometry data were analysed using MaxQuant software (v2.4.7.0) and utilising the Andromeda search engine.^56^ The MS/MS spectra were aligned to the Uniprot Homo sapiens database and the corresponding influenza virus Uniprot protein database. Known mass spectrometry contaminants and reversed sequences were also included. The search was performed with trypsin selected as the specific enzyme and a maximum of 2 missed cleavage events allowed. The MS accuracy was set to 10 ppm, then 0.05 Da for the MS/MS. The maximum peptide charge was set to seven and seven amino acids were required for the minimum peptide length. One unique peptide to the protein group was required to call the presence of a specific protein, while a false-discovery rate (FDR) of 1% was set. Statistical analysis to ascertain which proteins are enriched in one set of biological samples over another was performed using the intensities quantified by MaxQuant on Perseus software (v2.0.11).^57^ First, any potential contaminants, reverse hits or proteins that were not well identified (above 1% FDR) were removed. After filtering, the intensities were log-transformed (log2) and entries absent from at least 70% of the replicates for a given condition were removed. Missing values were replaced from the normal distribution. The data were then normalised by subtracting the most frequent value and enrichments between conditions were calculated. Statistical significance was calculated by way of an unpaired two-tailed Student’s t-test for each protein group. These values were then log-transformed (-log10) and the resulting p values and enrichments were graphed as volcano plots. The cutoff for statistical significance was set to p < 0.05. Analysed data is included in Supplementary Tables S1-S4.

### siRNA knockdowns

For all siRNA knockdowns, siRNAs (TriFECTa DsiRNA Kit, IDT) targeting the specified mRNA, or a non-specific control siRNA were used. These siRNA sequences are listed in Supplementary Table S6. A final concentration between 20nM and 2nM total siRNA was used in each condition according to the experiment, and siRNAs were all transfected using Lipofectamine RNAiMAX (Invitrogen; 13778075) following the manufacturer’s instructions. A549 cells were seeded at the time of transfection and incubated for 24h before the media was changed to fresh growth media. Cells were then incubated for an additional 24h before further processing.

### IAV multicycle titres

For multicycle IAV infections, A549 cells were seeded and transfected with siRNA simultaneously (20nM), as described above. At 48 h post-transfection cells were infected with WSN_PA-mNeon at an MOI 0.01 for 48h in IAV growth media, as described above, supplemented with 1:10000 TPCK trypsin. Supernatants were then titred for influenza virus on MDCK cells by plaque assay.

For the titrations of other IAV strains, they were carried out as above, but for Brisbane18 and Shanghai90, an MOI of 0.1 was used.

### IAV qPCR analysis

A549 cells previously transfected with siRNAs for 48h, were infected with A/WSN/33 virus at an MOI 3 PFU/cell. At 8 hpi RNA was extracted from cells using TRIzol (Invitrogen; 15596026). RNA was isolated and precipitated following the manufacturer’s instructions and 200ng was reverse transcribed into cDNA using the ABI cDNA synthesis kit (Applied Biosystems; 4368814). For cellular targets GAPDH and MYOF, RT was performed using an oligo dT, while for viral NP mRNA and NP vRNA specific primers were used, as described previously.^58^ These primers are listed in Table S7 alongside the primers used for qPCR amplification. All qPCR experiments were performed using SYBR Select Master Mix (Applied Biosystems; 4472908) following the manufacturer’s instructions. All qPCR data were normalised to GAPDH and quantified using the ΔΔCT method in relation to the siRNA NSC transfected cells.

### Immunofluorescence

Approximately 1.5*10^5 A549 (or A549_Rab11-mCherry) cells were seeded on glass coverslips (thickness #1.5) for 24h prior to treatment. For experiments involving knockdown, siRNAs were transfected as stated above during seeding and cells were cultured for 48h before further treatment.

Cells were then infected/treated according to the experiment, then fixed in either PFA 4% for 10’ followed by permeabilization by Triton-X-100 0.2% for 20’, or ice-cold methanol for 5’. The latter proved much better for the visualisation of MYOF and was used predominantly in experiments with wild-type A549s. Samples were then blocked in 3% BSA for 30’, followed by overnight incubation at 4°C with the primary antibodies diluted in 3% BSA. The next day, cells were washed and incubated for 1h with the secondary antibodies diluted in 3% BSA, then counterstained with Hoechst (Abcam; ab228551, 1:4000) before being mounted on slides using ProLong Diamond Antifade Mount (Thermo Fisher Scientific; P36961). Antibodies and dilutions are listed in Table S5.

### smiFISH

For smiFISH experiments, we employed the protocol described in Tsanov et al., using primary probes we designed (Supplementary Table S8, IDT oligo pool), annealed to the published imager strands (IDT Cy3- or Cy5-conjugated oligos).^59^ After probe hybridisation and subsequent washes, we proceeded with blocking and IF as described above.

### Lentiviral overexpression cell line generation

The Rab11-mCherry, EHD1-eGFP and EHD2-eGFP cell lines were generated by transduction with third-generation lentiviral vectors. Briefly, the producer 293T cell line was transfected with a 3:2:1 ratio of d8.74:pMD2.G:lentiviral vector, using PEI according to the manufacturer’s protocol. The lentiviral vector expressing mCherry-tagged Rab11 was pHR-FKBP:mCherry-Rab11a (Addgene plasmid # 72902; http://n2t.net/addgene:72902; RRID:Addgene_72902). The lentiviral vectors expressing eGFP-tagged EHD1 and EHD2 were cloned into the lentiviral vector pLCE and generated by PCR using a synthetic gene block as the template for EHD1 and the EHD2-mEGFP plasmid (Addgene plasmid #45932; http://n2t.net/addgene:45932; RRID:Addgene_45932) as the template for EHD2. The day after, the transfection media was exchanged for fresh media and incubated at 37°C for 72h. Then, the infectious supernatant was filtered with a 0.45 µm filter and overlaid onto fresh A549 cells. After 72h, Rab11a-mCherry cells were subjected to clonal isolation by dilution, cultured and screened for localisation of fluorescence signal, while EHD1-eGFP and EHD2-eGFP cell lines remained polyclonal for infection experiments.

### Transferrin uptake assay

For the transferrin uptake experiments, approximately 7.5*10^4 A549 cells per condition were seeded and transfected with the appropriate siRNA as described above. 48h post-transfection, the cells were serum starved (in serum-free media) for 30’ at 4°C, then incubated with fluorescently labelled transferrin at a concentration of 7.8 µg/ml for 30’ at 37°C. Cells were then washed with ice-cold PBS and fixed with 4% PFA for 10’, before further processing for IF.

Human holo-transferrin (Merck; T0665) was conjugated to Atto 647N NHS ester (Merck; 18373), in sodium carbonate buffer, according to manufacturer’s instructions. Excess dye was removed using Zeba™ Spin Desalting Columns (Thermo Fisher Scientific; 89877).

### Image analysis

Coverslips were imaged with a Leica Stellaris SP8 confocal microscope with a 100x objective with 1.4 NA. Detectors were set according to the dyes used, and each experiment had bleed-through controls to ensure signal specificity.

All analysis was carried out in FIJI using custom macros.

Briefly, for colocalisation analysis, the BIOP implementation of the JACoP plugin was used to quantify single-cell level pairwise correlation between signals in single z-slices after nuclear masking.^62^ At least 10 cells/condition per replicate were analysed across three biological replicates.

For the relocalisation experiments and the transferrin uptake assay, whole z-stacks were analysed by maximum projection, followed by automatic entropy-based thresholding of the signal of interest, and measurement of the area of signal foci. The resulting datasets were filtered for outliers in GraphPad Prism using the ROUT method and Q=1%. Roughly 30 cells/condition per replicate were analysed across three biological replicates.

### Nucleozin treatment

Nucleozin (MedChem Express; HY-50001) was reconstituted and stored in DMSO at a concentration of 5 mM. Mock/infected cells were treated with 1 µM nucleozin in infection media for 2h prior to fixation.

### RSV/SeV infection and titre

Isolation and characterisation of the clinical isolate RSV-BT2a was previously described.^60^ RSV A2/mKate2 was rescued by the Power Group from an infectious clone and helper plasmids kindly provided by Dr Martin Moore (Emroy University, Atlanta, USA).

Sendai_eGFP was rescued as previously described.^61^

For the titration experiments, approximately 1.5*10^5 A549 cells per condition were seeded and transfected with the appropriate siRNA as stated above. 48h post-transfection, cells were infected with MOI 0.1 RSV_mKate2, or MOI 0.1 SeV_eGFP, for 72h. Supernatants were then collected and titred. Hep-2 cells were used for RSV titrations, as previously described.^52^

For imaging, A549_Rab11-mCherry-HA cells were infected with MOI 1 RSV wild type BT2a or SeV_eGFP for 72h and processed for smiFISH and IF as described above.

### Statistical analysis

Unless otherwise stated, all experiments were performed as 3 independent biological replicates. When bar graphs are used to represent the data, each individual replicate can be observed as a point overlaid on the bar. Significance was calculated by performing two-way ANOVA with * indicating p<0.05, ** indicating p<0.01, *** indicating p<0.001, and **** indicating p<0.0001.

## Notes

### Competing Interest Statement

The authors have declared no competing interest.

