## Supplementary Information for "Myoferlin is an essential component of late-stage vRNP trafficking vesicles for enveloped RNA viruses"

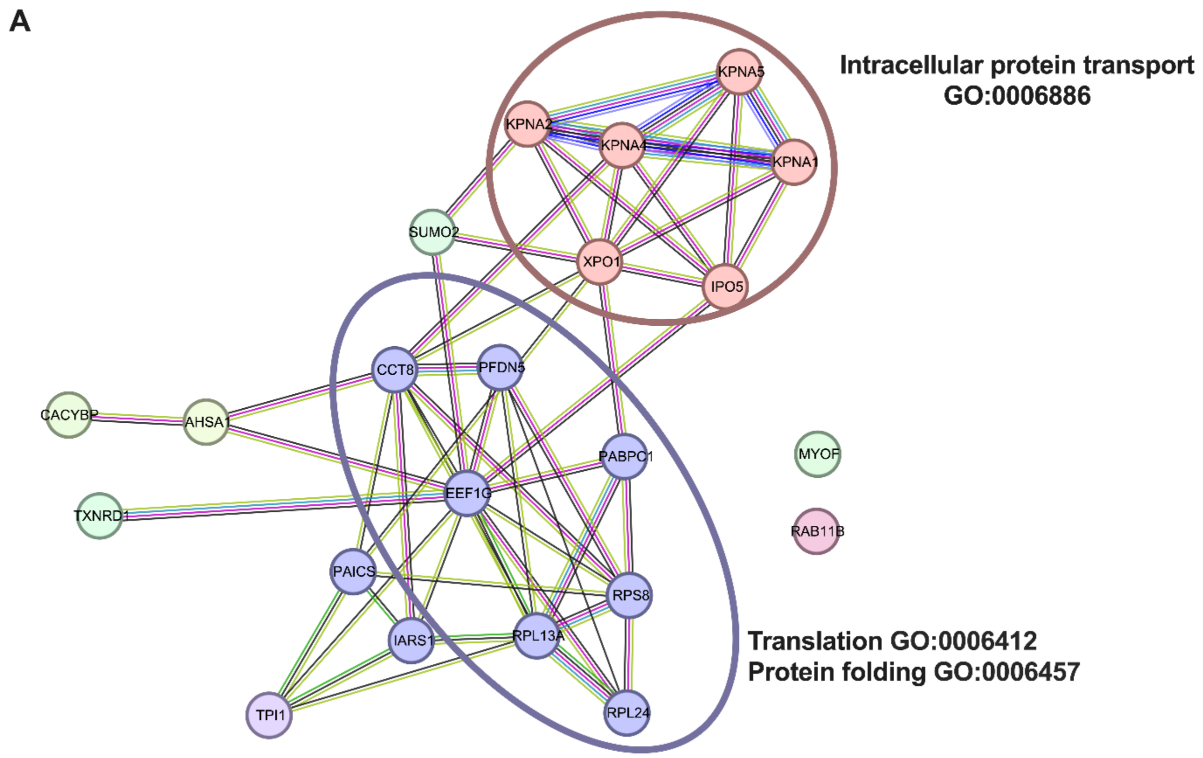

**Supplementary Figure 1 – STRING analysis of interactome at 16 hpi.**

(A) STRING analysis of all host proteins significantly enriched ($p<0.05$) in the 16 hpi PA-Flag interactome visualised in Figure 1E. Clusters of proteins that are part of the GO biological processes ‘Intracellular protein transport’ and ‘Translation and Protein folding’ are encircled and labelled.

A

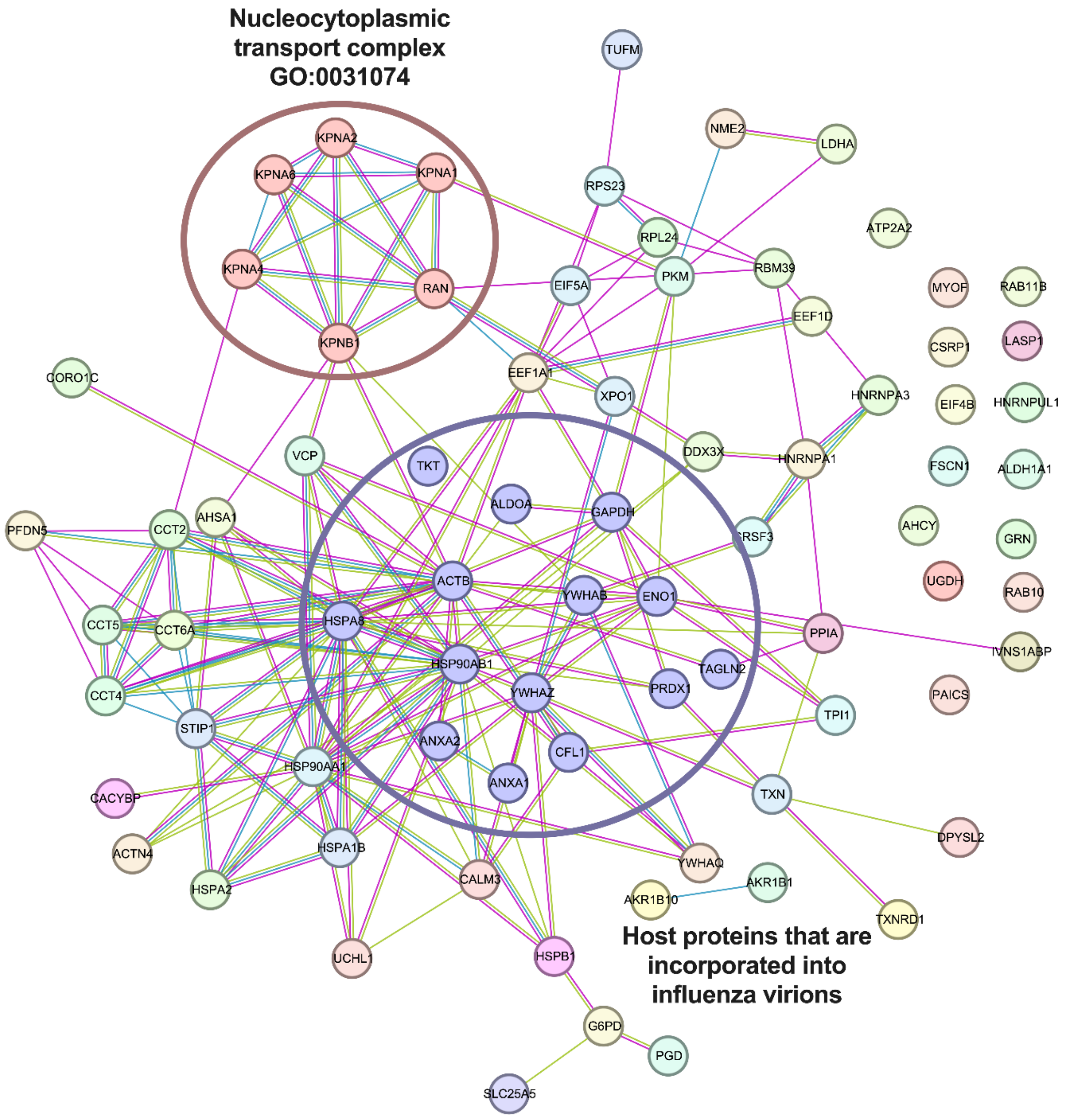

**Supplementary Figure 2 – STRING analysis of interactors enriched at 16 over 6 hpi.**

(A) STRING analysis of all host proteins significantly enriched (p<0.05 and >3-fold) in the 16 hpi PA-Flag interactome when compared to the 6 hpi interactome, visualised on the right side of the volcano plot in Figure 1G. A cluster of proteins involved in the GO cellular component of the 'Nucleocytoplasmic transport complex' and a different subset of proteins previously established as being incorporated into influenza A virus virions are encircled and labelled.[20]

**A**

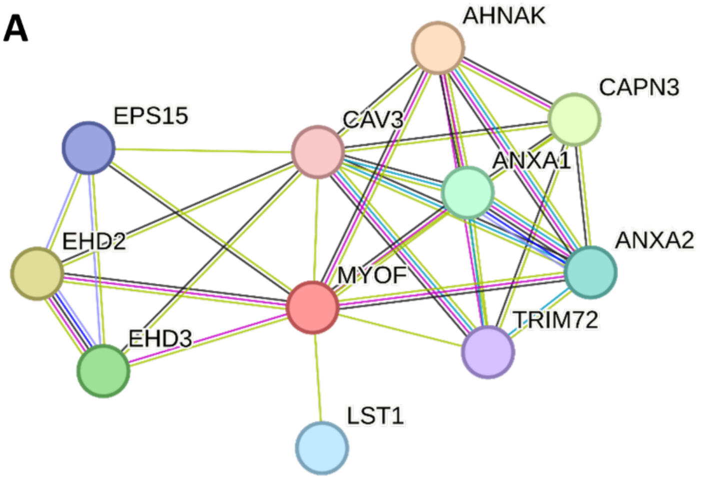

**B**

| #term ID | term description | strength | FDR | matching proteins in your network (labels) |
|---|---|---|---|---|
| GO:1905686 | Positive regulation of plasma membrane repair | 3.08 | 0.0103 | ANXA2,AHNAK |
| GO:0001765 | Membrane raft assembly | 2.6 | 0.0203 | CAV3,ANXA2 |
| GO:0031340 | Positive regulation of vesicle fusion | 2.6 | 0.0203 | ANXA2,ANXA1 |
| GO:0033292 | T-tubule organization | 2.6 | 0.0203 | CAV3,MYOF |
| GO:0001778 | Plasma membrane repair | 2.28 | 0.009 | TRIM72,CAV3,MYOF |
| GO:0032456 | Endocytic recycling | 1.91 | 0.0137 | EHD2,EHD3,EPS15 |
| GO:1901019 | Regulation of calcium ion transmembrane transporter activity | 1.74 | 0.0212 | EHD3,CAV3,AHNAK |
| GO:0007009 | Plasma membrane organization | 1.69 | 0.009 | EHD2,TRIM72,CAV3,MYOF |
| GO:1903169 | Regulation of calcium ion transmembrane transport | 1.59 | 0.0103 | EHD3,CAV3,AHNAK,CAPN3 |
| GO:0051668 | Localization within membrane | 1.22 | 0.0137 | EHD2,EHD3,CAV3,EPS15,CAPN3 |
| GO:0006897 | Endocytosis | 1.2 | 0.0493 | EHD2,EHD3,CAV3,EPS15 |
| GO:0051050 | Positive regulation of transport | 1.07 | 0.0103 | EHD2,EHD3,CAV3,ANXA2,ANXA1,CAPN3 |
| GO:0061024 | Membrane organization | 1.06 | 0.0311 | EHD2,TRIM72,CAV3,ANXA2,MYOF |
| GO:0016192 | Vesicle-mediated transport | 0.98 | 0.009 | EHD2,TRIM72,EHD3,CAV3,ANXA2,EPS15,ANXA1 |
| GO:0051049 | Regulation of transport | 0.85 | 0.0165 | EHD2,EHD3,CAV3,ANXA2,ANXA1,AHNAK,CAPN3 |
| GO:0016043 | Cellular component organization | 0.52 | 0.0212 | EHD2,TRIM72,EHD3,CAV3,ANXA2,MYOF,EPS15,LST1,ANXA1,CAPN3 |

**Supplementary Figure 3 – STRING analysis of myoferlin interacting proteins**

(A) STRING database query of all known MYOF interactions. (B) GO Biological Process term enrichment for the network in A.

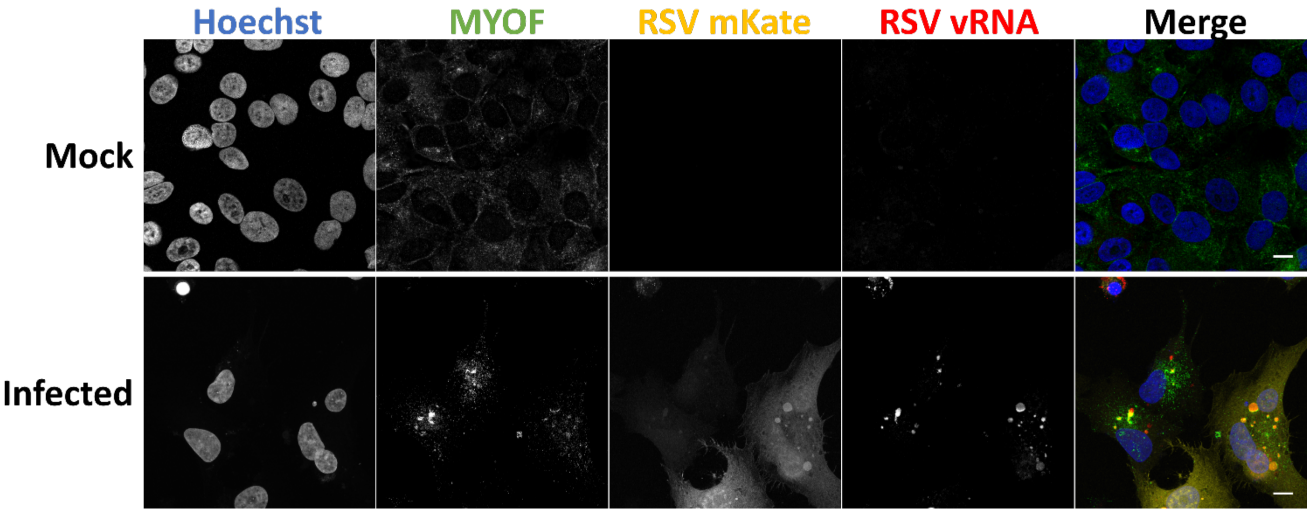

**Supplementary Figure 4 – IF and smiFISH of myoferlin associating with RSV_mKate vRNA**

A549 cells were infected with MOI1 of RSV_mKate2 for 72h, then stained for RSV vRNA and MYOF protein. Images are representative of two biological replicates. Scale bar 10 µm.

**Table S5 – Antibody dilutions**

| Target | Company | Species for secondary | Dilution |
|---|---|---|---|
| FLAG | Proteintech, 66008-3-ig | Ms | IF 1:500, WB 1:5000 |
| FLAG | Sigma, F1804 | Ms | IP 1:2 |
| PA | Thermo Fisher, Pa5-32223 | Rb | IF 1:500 |
| NP | Thermo Fisher, Pa5-32242 | Rb | IF 1:500, WB 1:5000 |
| NP | Abcam, Ab128193 | Ms | IF 1:500, WB 1:5000 |
| MYOF | Abcam, ab224091 | Rb | IF 1:200, WB 1:2500 |
| Actin | Proteintech, 66009-1-ig | Ms | WB 1:2500 |
| GAPDH | Proteintech, 60004-1-ig | Ms | WB 1:5000 |
| NS1 | Santa Cruz Biotechnology, Sc-130568 | Ms | WB 1:2500 |
| Rab11 | Invitrogen, 71-5300 | Rb | IF 1:100, WB 1:2500 |
| Ms, AlexaFluor 488 | Invitrogen, A32723 | / | IF 1:500 |
| Ms, AlexaFluor 647 | Invitrogen, A32728 | / | IF 1:500 |
| Rb, AlexaFluor 647 | Invitrogen, A32795 | / | IF 1:500 |
| Rb, AlexaFluor 488 | Invitrogen, A-11008 | / | IF 1:500 |
| Ms, HRP | Sigma, A9044 | / | WB 1:5000 |
| Rb, HRP | Sigma, A6154 | / | WB 1:5000 |

**Table S6 – siRNA sequences**

| Name | Sequence (5' → 3') |
|---|---|
| MYOF 1 | rGrCrU rCrUrA rArArA rUrCrA rGrGrG rArUrA rCrArA rGrGT A |
| MYOF 2 | rGrUrC rArUrU rGrArA rGrArC rCrGrA rGrArU rCrArC rUrAC A |
| PFDN5 1 | rArArC rArUrG rGrCrG rCrArG rUrCrU rArUrU rArArC rArUC A |
| PFDN5 2 | rArUrA rGrArU rUrUrU rCrUrA rArCrC rArArG rCrArG rArUG G |
| EEIF1G 1 | rUrUrA rCrUrU rGrArA rGrArC rGrArG rGrArC rUrUrU rUrCT G |
| EEIF1G 2 | rCrGrA rGrUrG rArCrA rUrUrG rGrCrU rGrArC rArUrC rArCA G |
| CCT8 1 | rGrUrA rCrUrC rArArG rArArU rCrArC rCrUrG rArUrG rUrUT T |
| CCT8 2 | rGrGrA rCrUrU rGrArA rCrArG rUrArU rGrCrU rArUrU rArAG A |
| PAICS 1 | rGrArA rGrUrG rGrArU rArUrC rArUrG rArGrU rCrArU rGrCT A |
| PAICS 2 | rGrArA rGrArU rGrUrC rArUrU rGrUrA rCrArC rUrUrU rArUT T |
| CACYBP 1 | rGrCrA rGrArA rCrUrU rCrUrU rGrArU rArArU rGrArA rArAA C |
| CACYBP 2 | rGrGrU rGrUrU rUrCrU rArUrC rUrArG rArUrU rUrArA rArUA T |
| RAB11 1 | rCrUrA rGrUrG rArCrC rUrUrA rCrArU rCrArG rUrGrA rUrCT T |
| RAB11 2 | rArGrA rUrCrA rUrGrC rUrGrA rUrArG rUrArA rCrArU rUrGT T |
| PABPC1 1 | rCrUrArGrGrGrCrArArArArGrArArUrUrCrArCrCrArArUGT |
| PABPC1 2 | rGrCrCrArGrGrUrCrUrArGrCrArArArCrArUrArArUrGrCTA |

**Table S7 – Primer sequences**

| | |
|---|---|
| WSN_NP_mRNA.RT | 5' CCAGATCGTTCGAGTCGTTTTTTTTTTTTTTTTTCTTTAATTGTC 3' |
| WSN_NP_vRNA.RT | 5' GGCCGTCATGGTGGCGAATGAATGGACGGAGAACAAGGATTGC 3' |
| Influenza_mRNA.R | 5' CCAGATCGTTCGAGTCGT 3' |
| Influenza_vRNA.R | 5' GGCCGTCATGGTGGCGAAT 3' |
| WSN_NP_mRNA.F | 5' CGATCGTGCCCTCCTTTG 3' |
| WSN_NP_vRNA.F | 5' CTCAATATGAGTGCAGACCGTGCT 3' |
| GAPDH.F | 5' TGGGTGTGAACCATGAGAAG 3' |

| | |
|---|---|
| GAPDH.R | 5’ GATGGCATGGACTGTGGTC 3’ |
| MYOF.F | 5’ CCCAGGCTGATCATTCAGAT 3’ |
| MYOF.R | 5’ CAAAGAGGGAGGCTGTCTTG 3’ |

**Table S8 – smiFISH probes**

| **Probe Name** | **Sequence** |
|---|---|
| **RSV** | |
| RSV_gRNA_smiFISH-01 | ctgcgaatatattcccagtaCCTCCTAAGTTTCGAGCTGGACTCAGTG |
| RSV_gRNA_smiFISH-02 | ggatcggaggtttacttagtCCTCCTAAGTTTCGAGCTGGACTCAGTG |
| RSV_gRNA_smiFISH-03 | gtatgcttaggcagtaagttCCTCCTAAGTTTCGAGCTGGACTCAGTG |
| RSV_gRNA_smiFISH-04 | caagtactgttcctcagttaCCTCCTAAGTTTCGAGCTGGACTCAGTG |
| RSV_gRNA_smiFISH-05 | ctgctacagatgcaaccaacCCTCCTAAGTTTCGAGCTGGACTCAGTG |
| RSV_gRNA_smiFISH-06 | gcgtacagtagtggaacttcCCTCCTAAGTTTCGAGCTGGACTCAGTG |
| RSV_gRNA_smiFISH-07 | atcactttactgcatgcttcCCTCCTAAGTTTCGAGCTGGACTCAGTG |
| RSV_gRNA_smiFISH-08 | atttattccctatggttgtgCCTCCTAAGTTTCGAGCTGGACTCAGTG |
| RSV_gRNA_smiFISH-09 | gcaaagcaaagctggagtgtCCTCCTAAGTTTCGAGCTGGACTCAGTG |
| RSV_gRNA_smiFISH-10 | agacctatctcttgtgttttCCTCCTAAGTTTCGAGCTGGACTCAGTG |
| RSV_gRNA_smiFISH-11 | gctggacattggattctgatCCTCCTAAGTTTCGAGCTGGACTCAGTG |
| RSV_gRNA_smiFISH-12 | gatgaaacctcccatattcaCCTCCTAAGTTTCGAGCTGGACTCAGTG |
| RSV_gRNA_smiFISH-13 | agctttggccttagtttaatCCTCCTAAGTTTCGAGCTGGACTCAGTG |
| RSV_gRNA_smiFISH-14 | gacactagccctattaatcgCCTCCTAAGTTTCGAGCTGGACTCAGTG |
| RSV_gRNA_smiFISH-15 | agtcagtagtagaccatgtgCCTCCTAAGTTTCGAGCTGGACTCAGTG |
| RSV_gRNA_smiFISH-16 | agaactcagcataggaacccCCTCCTAAGTTTCGAGCTGGACTCAGTG |
| RSV_gRNA_smiFISH-17 | attggattgggtgtatgcatCCTCCTAAGTTTCGAGCTGGACTCAGTG |
| RSV_gRNA_smiFISH-18 | cacccagtatcatgtatacaCCTCCTAAGTTTCGAGCTGGACTCAGTG |
| RSV_gRNA_smiFISH-19 | ggatacttccattggattgtCCTCCTAAGTTTCGAGCTGGACTCAGTG |
| RSV_gRNA_smiFISH-20 | gccactgagatgatgaggaaCCTCCTAAGTTTCGAGCTGGACTCAGTG |

| | |
|---|---|
| RSV_gRNA_smiFISH-21 | gaacctacatatcctcatggCCTCCTAAGTTTCGAGCTGGACTCAGTG |
| RSV_gRNA_smiFISH-22 | cagaggttttgagtacagctCCTCCTAAGTTTCGAGCTGGACTCAGTG |
| RSV_gRNA_smiFISH-23 | agggtctgagagacaagctaCCTCCTAAGTTTCGAGCTGGACTCAGTG |
| RSV_gRNA_smiFISH-24 | tgatgagagatcctcaagctCCTCCTAAGTTTCGAGCTGGACTCAGTG |
| RSV_gRNA_smiFISH-25 | accctaatgctgaattcgtaCCTCCTAAGTTTCGAGCTGGACTCAGTG |
| RSV_gRNA_smiFISH-26 | agagtgggaccgtggataaaCCTCCTAAGTTTCGAGCTGGACTCAGTG |
| RSV_gRNA_smiFISH-27 | tgtatattacccagctagtaCCTCCTAAGTTTCGAGCTGGACTCAGTG |
| RSV_gRNA_smiFISH-28 | caggcataggccacaaattaCCTCCTAAGTTTCGAGCTGGACTCAGTG |
| RSV_gRNA_smiFISH-29 | gctcaagcagattatttgctCCTCCTAAGTTTCGAGCTGGACTCAGTG |
| RSV_gRNA_smiFISH-30 | atcagactcatggaaggtcaCCTCCTAAGTTTCGAGCTGGACTCAGTG |
| RSV_gRNA_smiFISH-31 | actgtggaccatagaagctaCCTCCTAAGTTTCGAGCTGGACTCAGTG |
| RSV_gRNA_smiFISH-32 | tctctattttcctggttacaCCTCCTAAGTTTCGAGCTGGACTCAGTG |
| RSV_gRNA_smiFISH-33 | gtgctctatcatcacagatcCCTCCTAAGTTTCGAGCTGGACTCAGTG |
| RSV_gRNA_smiFISH-34 | tacatgccatcacacatacaCCTCCTAAGTTTCGAGCTGGACTCAGTG |
| RSV_gRNA_smiFISH-35 | atggactagtttccctagaaCCTCCTAAGTTTCGAGCTGGACTCAGTG |
| RSV_gRNA_smiFISH-36 | ggactacgtttctatcgtgaCCTCCTAAGTTTCGAGCTGGACTCAGTG |
| RSV_gRNA_smiFISH-37 | acttatccttctttgttggaCCTCCTAAGTTTCGAGCTGGACTCAGTG |
| RSV_gRNA_smiFISH-38 | atgctattgttttacccttaCCTCCTAAGTTTCGAGCTGGACTCAGTG |
| RSV_gRNA_smiFISH-39 | ttacaacagatggcctacttCCTCCTAAGTTTCGAGCTGGACTCAGTG |
| RSV_gRNA_smiFISH-40 | aagacaagccatggatgctgCCTCCTAAGTTTCGAGCTGGACTCAGTG |
| RSV_gRNA_smiFISH-41 | tatttggacacccaatggtaCCTCCTAAGTTTCGAGCTGGACTCAGTG |
| RSV_gRNA_smiFISH-42 | cttgcaggtgacaataacctCCTCCTAAGTTTCGAGCTGGACTCAGTG |
| RSV_gRNA_smiFISH-43 | gctatttcacaatgaggggtCCTCCTAAGTTTCGAGCTGGACTCAGTG |
| RSV_gRNA_smiFISH-44 | aggcttaagatgcggattcaCCTCCTAAGTTTCGAGCTGGACTCAGTG |
| RSV_gRNA_smiFISH-45 | tcctccatcatggttaatacCCTCCTAAGTTTCGAGCTGGACTCAGTG |
| RSV_gRNA_smiFISH-46 | gtcagaacagattgctaccaCCTCCTAAGTTTCGAGCTGGACTCAGTG |
| RSV_gRNA_smiFISH-47 | attcaatggtccttatctcaCCTCCTAAGTTTCGAGCTGGACTCAGTG |

| | |
|---|---|
| RSV_gRNA_smiFISH-48 | ggtgttatctctttctcagaCCTCCTAAGTTTCGAGCTGGACTCAGTG |
| **Sendai** | |
| Sendai_gRNA_smiFISH-01 | cttggcagagatatctagggCCTCCTAAGTTTCGAGCTGGACTCAGTG |
| Sendai_gRNA_smiFISH-02 | acgattctggaatgcaggtgCCTCCTAAGTTTCGAGCTGGACTCAGTG |
| Sendai_gRNA_smiFISH-03 | tcaaggatgtggaccttgagCCTCCTAAGTTTCGAGCTGGACTCAGTG |
| Sendai_gRNA_smiFISH-04 | aagccactgatattgcacttCCTCCTAAGTTTCGAGCTGGACTCAGTG |
| Sendai_gRNA_smiFISH-05 | gttgaaagatacggcaacccCCTCCTAAGTTTCGAGCTGGACTCAGTG |
| Sendai_gRNA_smiFISH-06 | tgctctaagacacgtcatgtCCTCCTAAGTTTCGAGCTGGACTCAGTG |
| Sendai_gRNA_smiFISH-07 | tgactactctttctgtctcaCCTCCTAAGTTTCGAGCTGGACTCAGTG |
| Sendai_gRNA_smiFISH-08 | tcataggcttcaagtttcggCCTCCTAAGTTTCGAGCTGGACTCAGTG |
| Sendai_gRNA_smiFISH-09 | ctatacgagtgtcatgcagtCCTCCTAAGTTTCGAGCTGGACTCAGTG |
| Sendai_gRNA_smiFISH-10 | tgacagggtatatcctaaccCCTCCTAAGTTTCGAGCTGGACTCAGTG |
| Sendai_gRNA_smiFISH-11 | taagatcaggtctctgggtaCCTCCTAAGTTTCGAGCTGGACTCAGTG |
| Sendai_gRNA_smiFISH-12 | acatccacaatatctctcagCCTCCTAAGTTTCGAGCTGGACTCAGTG |
| Sendai_gRNA_smiFISH-13 | tcagaagagcctgaatacctCCTCCTAAGTTTCGAGCTGGACTCAGTG |
| Sendai_gRNA_smiFISH-14 | catataccacgacatcgtgtCCTCCTAAGTTTCGAGCTGGACTCAGTG |
| Sendai_gRNA_smiFISH-15 | actgatgcttatccattgtcCCTCCTAAGTTTCGAGCTGGACTCAGTG |
| Sendai_gRNA_smiFISH-16 | gtattcctaacccatgacaaCCTCCTAAGTTTCGAGCTGGACTCAGTG |
| Sendai_gRNA_smiFISH-17 | atttcggaaccatcggagagCCTCCTAAGTTTCGAGCTGGACTCAGTG |
| Sendai_gRNA_smiFISH-18 | taatggatgtgaatccccatCCTCCTAAGTTTCGAGCTGGACTCAGTG |
| Sendai_gRNA_smiFISH-19 | ccttgcagaccttgcattgcCCTCCTAAGTTTCGAGCTGGACTCAGTG |
| Sendai_gRNA_smiFISH-20 | gtaagtatgacctgtatcctCCTCCTAAGTTTCGAGCTGGACTCAGTG |
| Sendai_gRNA_smiFISH-21 | gaggacagcaagacagtgttCCTCCTAAGTTTCGAGCTGGACTCAGTG |
| Sendai_gRNA_smiFISH-22 | agcattacagtgtaatccggCCTCCTAAGTTTCGAGCTGGACTCAGTG |
| Sendai_gRNA_smiFISH-23 | agctgcttgaaagcacttgaCCTCCTAAGTTTCGAGCTGGACTCAGTG |
| Sendai_gRNA_smiFISH-24 | agtattcccatctactttgaCCTCCTAAGTTTCGAGCTGGACTCAGTG |

| | |
|---|---|
| Sendai_gRNA_smiFISH-25 | ctgtgacaatgacccagatgCCTCCTAAGTTTCGAGCTGGACTCAGTG |
| Sendai_gRNA_smiFISH-26 | tctgtggatctattcatgcaCCTCCTAAGTTTCGAGCTGGACTCAGTG |
| Sendai_gRNA_smiFISH-27 | cctaacctctcaaatcaggaCCTCCTAAGTTTCGAGCTGGACTCAGTG |
| Sendai_gRNA_smiFISH-28 | tattttgctcaacgttcgggCCTCCTAAGTTTCGAGCTGGACTCAGTG |
| Sendai_gRNA_smiFISH-29 | cagagcaaccagtatctgtaCCTCCTAAGTTTCGAGCTGGACTCAGTG |
| Sendai_gRNA_smiFISH-30 | ccaaggtccacattagatttCCTCCTAAGTTTCGAGCTGGACTCAGTG |
| Sendai_gRNA_smiFISH-31 | cagattatgctaactgggctCCTCCTAAGTTTCGAGCTGGACTCAGTG |
| Sendai_gRNA_smiFISH-32 | cgtgcaagtcggttcataacCCTCCTAAGTTTCGAGCTGGACTCAGTG |
| Sendai_gRNA_smiFISH-33 | acctggacacgcttacaaacCCTCCTAAGTTTCGAGCTGGACTCAGTG |
| Sendai_gRNA_smiFISH-34 | gcagtacgagacactgactaCCTCCTAAGTTTCGAGCTGGACTCAGTG |
| Sendai_gRNA_smiFISH-35 | tcaccgagactagtggagaaCCTCCTAAGTTTCGAGCTGGACTCAGTG |
| Sendai_gRNA_smiFISH-36 | catcagagcggatctgttagCCTCCTAAGTTTCGAGCTGGACTCAGTG |
| Sendai_gRNA_smiFISH-37 | gttgtataagacctgtcagcCCTCCTAAGTTTCGAGCTGGACTCAGTG |
| Sendai_gRNA_smiFISH-38 | gatgtagggcacgagctaaaCCTCCTAAGTTTCGAGCTGGACTCAGTG |
| Sendai_gRNA_smiFISH-39 | gttcagggtgacaatcaagcCCTCCTAAGTTTCGAGCTGGACTCAGTG |
| Sendai_gRNA_smiFISH-40 | agagtactgcattgtttggtCCTCCTAAGTTTCGAGCTGGACTCAGTG |
| Sendai_gRNA_smiFISH-41 | gttaagttgcttcctcacaaCCTCCTAAGTTTCGAGCTGGACTCAGTG |
| Sendai_gRNA_smiFISH-42 | cttataagatgcgagccgtaCCTCCTAAGTTTCGAGCTGGACTCAGTG |
| Sendai_gRNA_smiFISH-43 | tccatggaacctctattgatCCTCCTAAGTTTCGAGCTGGACTCAGTG |
| Sendai_gRNA_smiFISH-44 | agcagacactattgtggagtCCTCCTAAGTTTCGAGCTGGACTCAGTG |
| Sendai_gRNA_smiFISH-45 | tgatcctgttatacctctacCCTCCTAAGTTTCGAGCTGGACTCAGTG |
| Sendai_gRNA_smiFISH-46 | gacaaaatccgatccgtcttCCTCCTAAGTTTCGAGCTGGACTCAGTG |
| Sendai_gRNA_smiFISH-47 | atacacgtttgaaccgtaccCCTCCTAAGTTTCGAGCTGGACTCAGTG |
| Sendai_gRNA_smiFISH-48 | cagactgaaggacgacagcaCCTCCTAAGTTTCGAGCTGGACTCAGTG |
| **WSN NA vRNA (seg6)** | |
| smiFISH_WSN_NA-vRNA-1 | gttcaccattgacaagtagtCCTCCTAAGTTTCGAGCTGGACTCAGTG |

| | |
|---|---|
| smiFISH_WSN_NA-vRNA-2 | tactgtagattggtcttggcCCTCCTAAGTTTCGAGCTGGACTCAGTG |
| smiFISH_WSN_NA-vRNA-3 | ggacgcaatctggactagtgCCTCCTAAGTTTCGAGCTGGACTCAGTG |
| smiFISH_WSN_NA-vRNA-4 | taatcagggggctacctgagCCTCCTAAGTTTCGAGCTGGACTCAGTG |
| smiFISH_WSN_NA-vRNA-5 | ctagactgtatgaggccttgCCTCCTAAGTTTCGAGCTGGACTCAGTG |
| smiFISH_WSN_NA-vRNA-6 | tcaacatcctgagctaacagCCTCCTAAGTTTCGAGCTGGACTCAGTG |
| smiFISH_WSN_NA-vRNA-7 | cagggtacagcggaagtttcCCTCCTAAGTTTCGAGCTGGACTCAGTG |
| smiFISH_WSN_NA-vRNA-8 | gttgtggcaatgactgatcgCCTCCTAAGTTTCGAGCTGGACTCAGTG |
| smiFISH_WSN_NA-vRNA-9 | agtaggttctctatgagacaCCTCCTAAGTTTCGAGCTGGACTCAGTG |
| smiFISH_WSN_NA-vRNA-10 | cctaatggatggacagagacCCTCCTAAGTTTCGAGCTGGACTCAGTG |
| smiFISH_WSN_NA-vRNA-11 | tgggtttgagatgatttgggCCTCCTAAGTTTCGAGCTGGACTCAGTG |
| smiFISH_WSN_NA-vRNA-12 | ggactaaaagtgacagttccCCTCCTAAGTTTCGAGCTGGACTCAGTG |
| smiFISH_WSN_NA-vRNA-13 | ggtaatggtgtttggataggCCTCCTAAGTTTCGAGCTGGACTCAGTG |
| smiFISH_WSN_NA-vRNA-14 | cggagtaaagggattttcatCCTCCTAAGTTTCGAGCTGGACTCAGTG |
| smiFISH_WSN_NA-vRNA-15 | cagtgtctgctgatggagcaCCTCCTAAGTTTCGAGCTGGACTCAGTG |
| smiFISH_WSN_NA-vRNA-16 | ccaaagatggaacaggcagcCCTCCTAAGTTTCGAGCTGGACTCAGTG |
| smiFISH_WSN_NA-vRNA-17 | taggatacatctgcagtgggCCTCCTAAGTTTCGAGCTGGACTCAGTG |
| smiFISH_WSN_NA-vRNA-18 | ccttcgaccaaaacctagatCCTCCTAAGTTTCGAGCTGGACTCAGTG |
| smiFISH_WSN_NA-vRNA-19 | ttaccctgataccggcaaagCCTCCTAAGTTTCGAGCTGGACTCAGTG |
| smiFISH_WSN_NA-vRNA-20 | ctcactacgaggaatgttccCCTCCTAAGTTTCGAGCTGGACTCAGTG |
| smiFISH_WSN_NA-vRNA-21 | caatagagttgaatgcacctCCTCCTAAGTTTCGAGCTGGACTCAGTG |
| smiFISH_WSN_NA-vRNA-22 | tcgagaaggggaaggttactCCTCCTAAGTTTCGAGCTGGACTCAGTG |
| smiFISH_WSN_NA-vRNA-23 | gctggcctcgtacaaaatttCCTCCTAAGTTTCGAGCTGGACTCAGTG |
| smiFISH_WSN_NA-vRNA-24 | ttaccataatgaccgatggcCCTCCTAAGTTTCGAGCTGGACTCAGTG |
| smiFISH_WSN_NA-vRNA-25 | gagtctgaatgtacctgtgtCCTCCTAAGTTTCGAGCTGGACTCAGTG |
| smiFISH_WSN_NA-vRNA-26 | aatgggctggctaacaatcgCCTCCTAAGTTTCGAGCTGGACTCAGTG |
| smiFISH_WSN_NA-vRNA-27 | cagcaagtgcatgtcatgatCCTCCTAAGTTTCGAGCTGGACTCAGTG |

| | |
|---|---|
| smiFISH_WSN_NA-vRNA-28 | cccgtacaattcaaggtttgCCTCCTAAGTTTCGAGCTGGACTCAGTG |
| smiFISH_WSN_NA-vRNA-29 | ttatagggccttaatgagctCCTCCTAAGTTTCGAGCTGGACTCAGTG |
| smiFISH_WSN_NA-vRNA-30 | ggggacctttaaggacagaaCCTCCTAAGTTTCGAGCTGGACTCAGTG |
| smiFISH_WSN_NA-vRNA-31 | cgccttactgaatgacaagcCCTCCTAAGTTTCGAGCTGGACTCAGTG |
| smiFISH_WSN_NA-vRNA-32 | ggaccttttttctgactcaaCCTCCTAAGTTTCGAGCTGGACTCAGTG |
| smiFISH_WSN_NA-vRNA-33 | tggttccaaaggagacgtttCCTCCTAAGTTTCGAGCTGGACTCAGTG |
| smiFISH_WSN_NA-vRNA-34 | gtgggctatacacagcaaagCCTCCTAAGTTTCGAGCTGGACTCAGTG |
| smiFISH_WSN_NA-vRNA-35 | cggcaattcatctctttgtcCCTCCTAAGTTTCGAGCTGGACTCAGTG |
| smiFISH_WSN_NA-vRNA-36 | ggactcaacttcagtgatatCCTCCTAAGTTTCGAGCTGGACTCAGTG |
| smiFISH_WSN_NA-vRNA-37 | cctataaagttgttgctgggCCTCCTAAGTTTCGAGCTGGACTCAGTG |
| smiFISH_WSN_NA-vRNA-38 | gaatatgcaaccaaggcagcCCTCCTAAGTTTCGAGCTGGACTCAGTG |
| smiFISH_WSN_NA-vRNA-39 | agccattcaattcaaaccggCCTCCTAAGTTTCGAGCTGGACTCAGTG |
| smiFISH_WSN_NA-vRNA-40 | agtcggaataattagcctaaCCTCCTAAGTTTCGAGCTGGACTCAGTG |
| smiFISH_WSN_NA-vRNA-41 | ccattggatcgatctgtatgCCTCCTAAGTTTCGAGCTGGACTCAGTG |
